# Adult newborn granule cells confer emotional-state-dependent plasticity in memory retrieval

**DOI:** 10.1101/2020.07.14.202481

**Authors:** Bo Lei, Bilin Kang, Wantong Lin, Haichao Chen, Yuejun Hao, Jian Ma, Songhai Shi, Yi Zhong

## Abstract

Achieving optimal behavior requires animals to flexibly retrieve prior knowledge. Here we show that adult newborn granule cells (anbGCs) mediate emotional-state-dependent plasticity of memory retrieval. We find that acute social reward (aSR) enhances memory retrieval by increasing the reactivation of engram cells, while acute social stress (aSS) weakens retrieval and reduces the reactivation. Such bidirectional regulation relies on the activation of distinct populations of anbGCs by aSR and aSS, triggering opposing modifications of dDG activity, which is sufficient to regulate and predict the performance of memory retrieval. Concordantly, in emotional disorder models, aSR-dependent memory plasticity is impaired, while the effect of aSS remains intact. Together, our data revealed that anbGCs mediate plasticity of memory retrieval, allowing animals to flexibly retrieve memory according to the current emotional state, and suggested the essential roles of anbGCs in translating emotional information to the regulation of memory expression.

## Introduction

Multiple hypotheses suggest that adult hippocampal neurogenesis could provide long-sought mechanisms for linking emotional experiences and memory (Aimone et al., 2009; Anacker and Hen, 2017; Leuner and Gould, 2010; Opendak and Gould, 2015). There are two lines of evidence supporting these hypotheses. First, adult neurogenesis is profoundly influenced by many behavioral factors (Praag et al., 2000), such as learning (Ambrogini et al., 2000), exercise (Van Praag et al., 1999), environmental enrichment (Kempermann et al., 1997), stress (Gould and Tanapat, 1999), and reward (Leuner et al., 2010; Ramirez et al., 2015). Second, ablating hippocampal neurogenesis or increasing the number of adult newborn granule cells (anbGCs) has been reported to affect many memory processes, including pattern separation (Nakashiba et al., 2012), adaptive forgetting (Akers et al., 2014), and memory flexibility (Burghardt et al., 2012). Consistent with these hypotheses, previous studies have revealed that chronic experience-triggered regulation of hippocampal neurogenesis is sufficient to affect memory performance. For instance, voluntary running can lead to adaptive forgetting by increasing hippocampal neurogenesis (Akers et al., 2014), while chronic stress can cause cognitive functional defects by decreasing hippocampal neurogenesis (Mckim et al., 2016). However, these effects are confined to the change in the number of anbGCs, which takes a long-time-interval to be effective and such mechanisms are inadequate to explain the impacts of pervasive daily experiences within a limited time window.

Extensive effort has been devoted to investigating how acute emotional experiences affect the memory process (Fernández and Morris, 2018; Joëls et al., 2011), enabling animals to flexibly retrieve memories according to their immediate emotional state. Acute stress can effectively impair memory retrieval in both animal models (Kim and Diamond, 2002; De Quervain et al., 1998; Tak et al., 2007) and humans (Kuhlmann, 2005; Smith et al., 2016) while the novel context, rewarding experiences, and natural emotional stress are capable of enhancing memory acquisition or maintenance (Hu et al., 2007; Singer and Frank, 2009; Takeuchi et al., 2016). However, how the hippocampus integrates the impacts of such diverse emotional experiences into memory performance is still poorly understood.

Here, we have used multiple approaches to examine how daily social experiences modulate memory retrieval and characterize the roles of anbGC activity in mediating such modulation. We found that acute social reward (aSR) and acute social stress (aSS) exert significant, differential impacts on the retrieval but not the formation of contextual fear memory by modifying the reactivation efficacy of engram cells. Such emotional-state-dependent bidirectional modulation of memory retrieval is mediated by the activation of distinct populations of anbGCs to trigger opposing regulation of dorsal dentate gyrus (dDG) activity.

## Results

### Acute social experiences regulate memory retrieval

Although efforts have been devoted to studying the effects of social experience on memory in humans (Kuhlmann, 2005; Takahashi et al., 2004), there are few behavioral paradigms available for assaying the impacts of acute daily social experiences on memory in animal models. Here, we designed two social interaction paradigms to assay the effects of aSR and aSS on memory.

After contextual fear conditioning (CFC), we exposed a conditioned male mouse to two female mice for 10 min as aSR or five grouped male mice as aSS (Figure 1A and see Methods). When acute social experiences were presented before the retrieval of recent memory (at 4 h or 24 h after CFC) (Figure 1A), aSR enhanced contextual fear memory retrieval significantly, while aSS reduced it (Figures 1B and 1C). We also used interaction with juvenile mice as aSR and social defeat as aSS, as reported in previous studies (Anacker et al., 2018; Golden et al., 2011; Mchenry et al., 2017; Mckim et al., 2016; Nardou et al., 2019), and observed similar regulatory effects on memory retrieval (Figure S1). There were no such effects when acute social experiences were presented immediately before CFC acquisition (Figures 1D-1F). In addition, the presentation of acute social experiences was effective only when given immediately or up to 0.5 h before memory recall and was ineffective if given 1 h, 1.5 h, or 2 h before memory recall (Figures 1G-1I). Thus, acute social experience paradigms regulated retrieval but not acquisition of recent memory.

**Figure 1.**
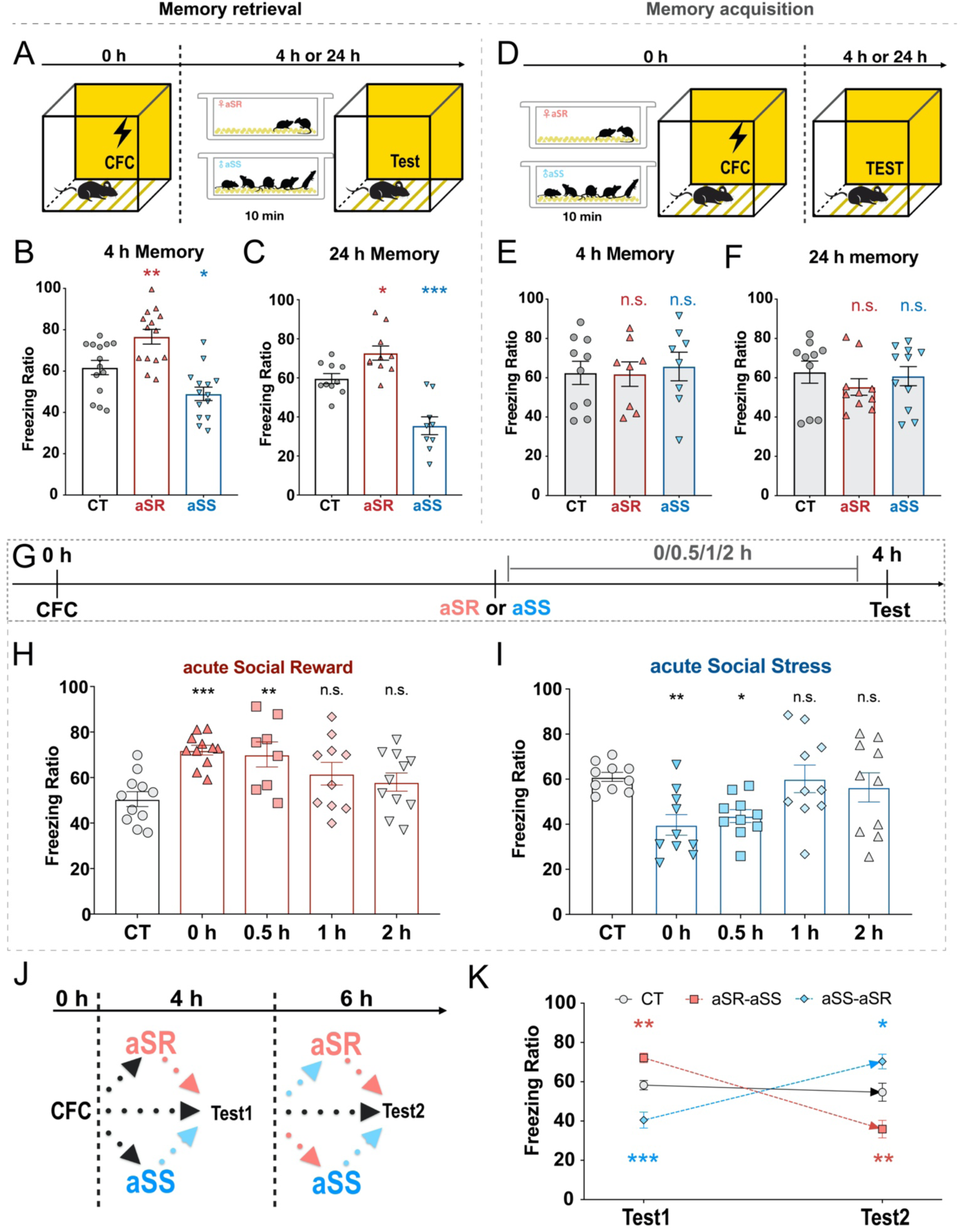
Acute social experience-dependent bidirectional regulation of retrieving recent contextual fear memory. **(A and D)** Experimental schedules of acute social experiences before memory retrieval (A) and memory acquisition (D). **(B and C)** Bidirectional regulation of acute social reward (aSR) and acute social stress (aSS) on retrieval of 4 h memory (B, n_CT_=14, n_aSR_=14, n_aSS_=14) and 24 h memory (C, n_CT_=10, n_aSR_=10, n_aSS_=9). **(E and F)** aSR and aSS before memory acquisition had no effect on 4 h memory (E, n_CT_=10, n_aSR_=8, n_aSS_=8) or 24 h memory (F, n_CT_=10, n_aSR_=10, n_aSS_=11). **(G)** Experimental schedule of the effective time window of acute social experiences. **(H)** 4 h memory affected by aSR at different time window (n_CT_=11, n_0h_=11, n_0.5h_=8, n_1h_=10, n_2h_=11). **(I)** 4 h memory affected by aSS at different time window (n_CT_=10, n_0h_=10, n_0.5h_=10, n_1h_=10, n_2h_=10). **(J)** Experimental schedule of two sequential retrieval tests. **(K)** Reversible regulations of acute social experiences on retrieval (n_CT_=16, n_aSR-aSS_=16, n_aSS-aSR_=16). *P < 0.05, **P < 0.01, ***P < 0.001; one-way analysis of variance (ANOVA) with Dunnett’s test. Graphs show means ± SEM. See also Figure S1.

We then performed two sequential retrieval tests to determine the dynamics of the regulation. The two tests were 2 h apart from each other, with every conditioned mouse being subjected to aSR immediately before one test and aSS immediately before another in counterbalanced order (Figure 1J). Recent contextual fear memory could be strengthened or weakened depending on the acute social experiences that occurred immediately prior to retrieval (Figure 1K). Overall, we observed temporally sensitive, dynamically regulated retrieval depending on acute social experiences.

### Adult hippocampal neurogenesis in dDG mediates effects of acute social experiences on memory retrieval

Since hippocampal neurogenesis is sensitive to diverse experiences (Ambrogini et al., 2000; Gould and Tanapat, 1999; Kempermann et al., 1997; Leuner and Gould, 2010; Leuner et al., 2010; Opendak and Gould, 2015; Praag et al., 2000; Van Praag et al., 1999; Ramirez et al., 2015) and shows a profound impact on memory processes (Akers et al., 2014; Anacker and Hen, 2017; Burghardt et al., 2012; Gu et al., 2012; Leuner and Gould, 2010; Nakashiba et al., 2012), anbGCs have been suggested to mediate the impacts of experiences on memory performance (Aimone et al., 2009; Anacker and Hen, 2017; Opendak and Gould, 2015). As a test of this possibility, we blocked neurogenesis to determine its effects on memory regulation by acute social experiences.

For this purpose, we irradiated the heads of mice with X-rays, which are thought to be able to significantly block neurogenesis (Kitamura et al., 2009; Monje et al., 2002) (Figures 2A-2C), and we found that the effects of acute social experiences on 4 h memory retrieval were eliminated in the irradiated mice (Figure 2D). To avoid these results being caused by the side effects of X-ray irradiation, such as induced neuroinflammation within the first month (Ben Abdallah et al., 2007), we conducted the behavior experiments 5 weeks after the irradiation treatment. We observed that the X-ray irradiated mice showed normal basal freezing level (before receiving foot-shock) in the CFC context, normal learning curve during CFC, and normal performance in anxiety-like behavior test (open field tests) (Figure S2), which was consistent with previous research showing X-ray irradiation-induced ablation of neurogenesis could not induce depression- or anxiety-like behaviors directly (Santarelli and Hen, 2003). To further confirm the effect of the ablation of adult neurogenesis, we next applied pharmacological treatment with temozolomide (TMZ) (Akers et al., 2014), which significantly decreases neurogenesis (Figure S3) with normal performance of basal freezing, memory acquisition and anxiety-like behavior (Figure S2). We observed impairment of acute social experience-based memory flexibility similar to the effects of X-ray irradiation (Figure S3).

**Figure 2.**
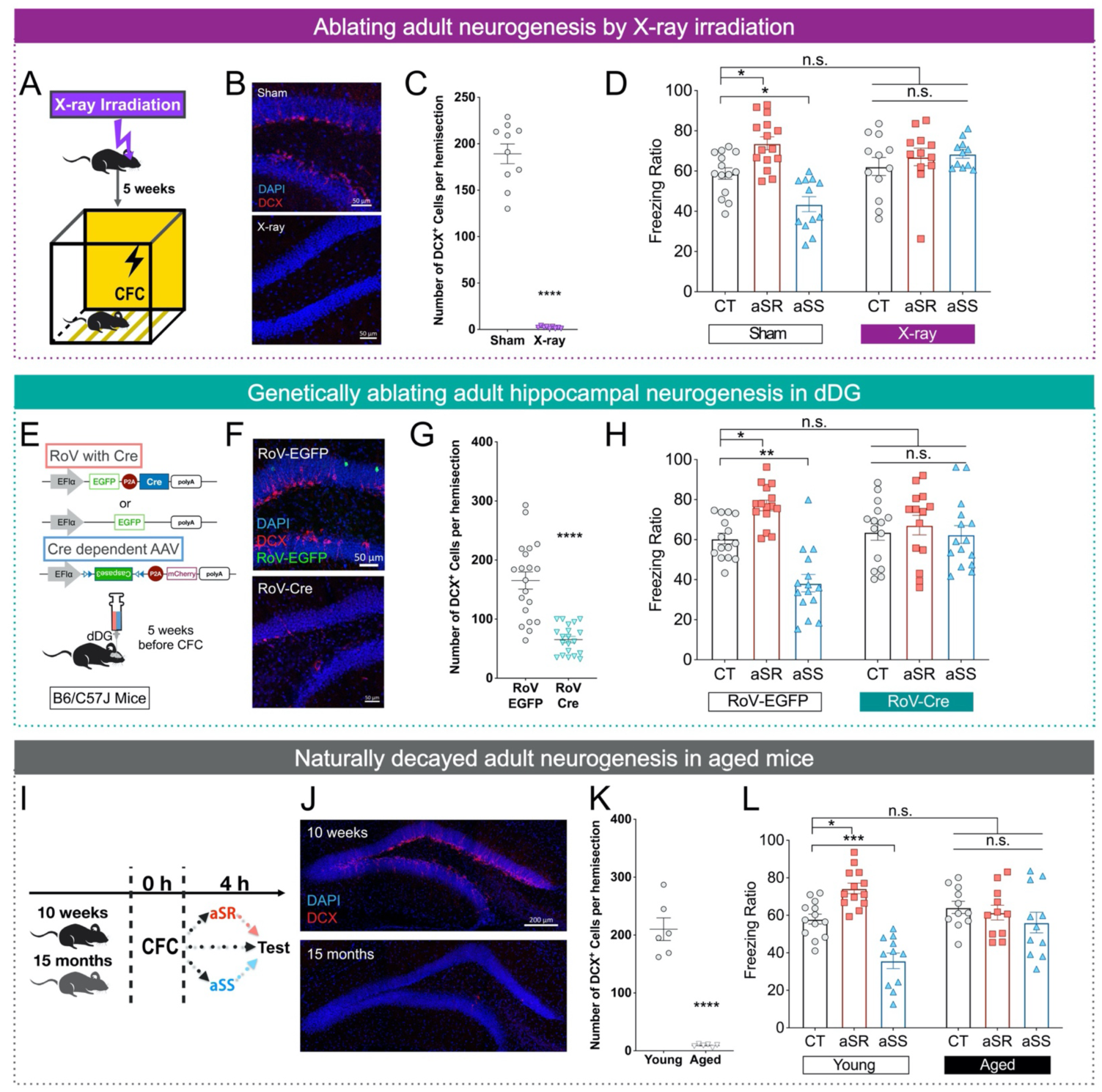
The effect of acute social experiences on memory retrieval depends on neurogenesis in the dorsal dentate gyrus (dDG). **(A)** Experimental schedule of adult newborn granule cells (anbGCs) ablation through X-ray irradiation. **(B)** Representative images of anbGCs (labelled by doublecortin, DCX) in the dDG. **(C)** Quantification of anbGCs in the dDG (n_Sham_=10, n_X-ray_=9). **(D)** 4 h memory in sham group (n_CT_=14, n_aSR_=15, n_aSS_=12) and X-ray group (n_CT_=12, n_aSR_=12, n_aSS_=11). **(E)** Experimental schedule of anbGCs ablation through caspase3 expression driven by double-virus system. **(F)** Representative images of anbGCs (DCX) in the dDG. **(G)** Quantification of anbGCs in the dDG (n_RoV-EGFP_=20, n_RoV-Cre_=20). **(H)** 4 h memory in RoV-EGFP group (n_CT_=15, n_aSR_=15, n_aSS_=15) and in RoV-Cre group (n_CT_=15, n_aSR_=14, n_aSS_=15). **(I)** Experimental schedule using aged mice. **(J)** Representative images of anbGCs (DCX) in the dDG. **(K)** Quantification of anbGCs in the dDG (n_Young_=6, n_Aged_=6). **(L)** 4 h memory in young mice (n_CT_=13, n_aSR_=13, n_aSS_=11) and in aged mice (n_CT_=11, n_aSR_=11, n_aSS_=11). *P < 0.05, **P < 0.01, ***P < 0.001, ****P < 0.0001; unpaired two-tailed t test (C, G and K), or two-way analysis of variance (ANOVA) with Tukey’s test (others). Graphs show means ± SEM. See also Figures S2-S4.

We sought to investigate whether neurogenesis in the dorsal dentate gyrus (dDG) is responsible for such memory flexibility, as the dDG has been reported to be significantly involved in memory processes (Anacker and Hen, 2017; Hainmueller and Bartos, 2020). For this purpose, we designed a double-virus system to specifically label and manipulate anbGCs in the dDG. In this system, we used a retrovirus (RoV) to target neural progenitors and their progeny (Fanselow and Dong, 2010; Tashiro et al., 2007) and express the cyclization recombination enzyme (Cre). Then, a Cre-dependent adeno-associated virus (AAV) drove the expression of caspase3 to induce apoptosis in neurons expressing Cre (Figure 2E). Five weeks after the injection of RoV and AAV in the dDG, the number of anbGCs (labelled with doublecortin, DCX) was significantly reduced in the dDG (Figures 2F and 2G), without affecting the number of anbGCs in the ventral dentate gyrus (Figure S4) and the performance of basal freezing, memory acquisition and anxiety-like behavior (Figure S2). With this manipulation, acute social experiences, whether aSR or aSS, had no influence on 4 h memory retrieval (Figure 2H).

Furthermore, aged mice (15 months old) with naturally reduced neurogenesis (Kuhn et al., 1996) (Figures 2I-2K) and normal performance of basal freezing, memory acquisition and anxiety-like behavior (Figure S2) also failed to show any effect of acute social experiences on retrieval (Figure 2L). The data presented indeed confirm the importance of adult hippocampal neurogenesis in mediating regulation of memory retrieval on the basis of acute social experience.

### Activation of anbGCs mediates effects of acute social experiences on memory retrieval

The generation and activation of anbGCs can affect cognitive function, such as memory forgetting (Akers et al., 2014), cognitive flexibility (Anacker and Hen, 2017; Burghardt et al., 2012), and pattern separation (Nakashiba et al., 2012). Since the acute social experience was presented only 10 min before retrieval, we suspected that anbGC activity would play a major role in the regulation of memory retrieval. To test whether anbGCs were activated during acute social experiences, conditioned mice were perfused 1.5 h after social interactions to label acute social experience-activated anbGCs by c-Fos and bromodeoxyuridine (BrdU) staining (Kee et al., 2007; Miller and Nowakowski, 1988) (Figures 3A and 3B). Compared to the control group, both acute social experience groups had increased proportions of c-Fos^+^ anbGCs in the overall anbGC population (Figure 3C), confirming the involvement of anbGC activation during acute social experiences. To validate the role of anbGC activation, we used the double-virus system to drive the expression of the optogenetic tool eNpHR3.0, which targeted anbGCs at a specific age (approximately 4 weeks old), in conjunction with RoV injected into the dDG at 4 weeks and AAV injected at 2 weeks before CFC (Figures 3D-3F). In the electrophysiological recording of brain slices, optical stimulation reliably silenced RoV-labelled anbGCs expressing eNpHR3.0 (Figure S5). Inhibiting anbGCs, i.e., stimulating eNpHR3.0 using a 589 nm laser, during acute social experiences blocked the regulatory effect of those experiences on memory retrieval (Figure 3G) without altering social interaction time (Figures S6E and S6F), suggesting an essential role of anbGC activation in the acute social experience-dependent regulation of memory retrieval.

**Figure 3.**
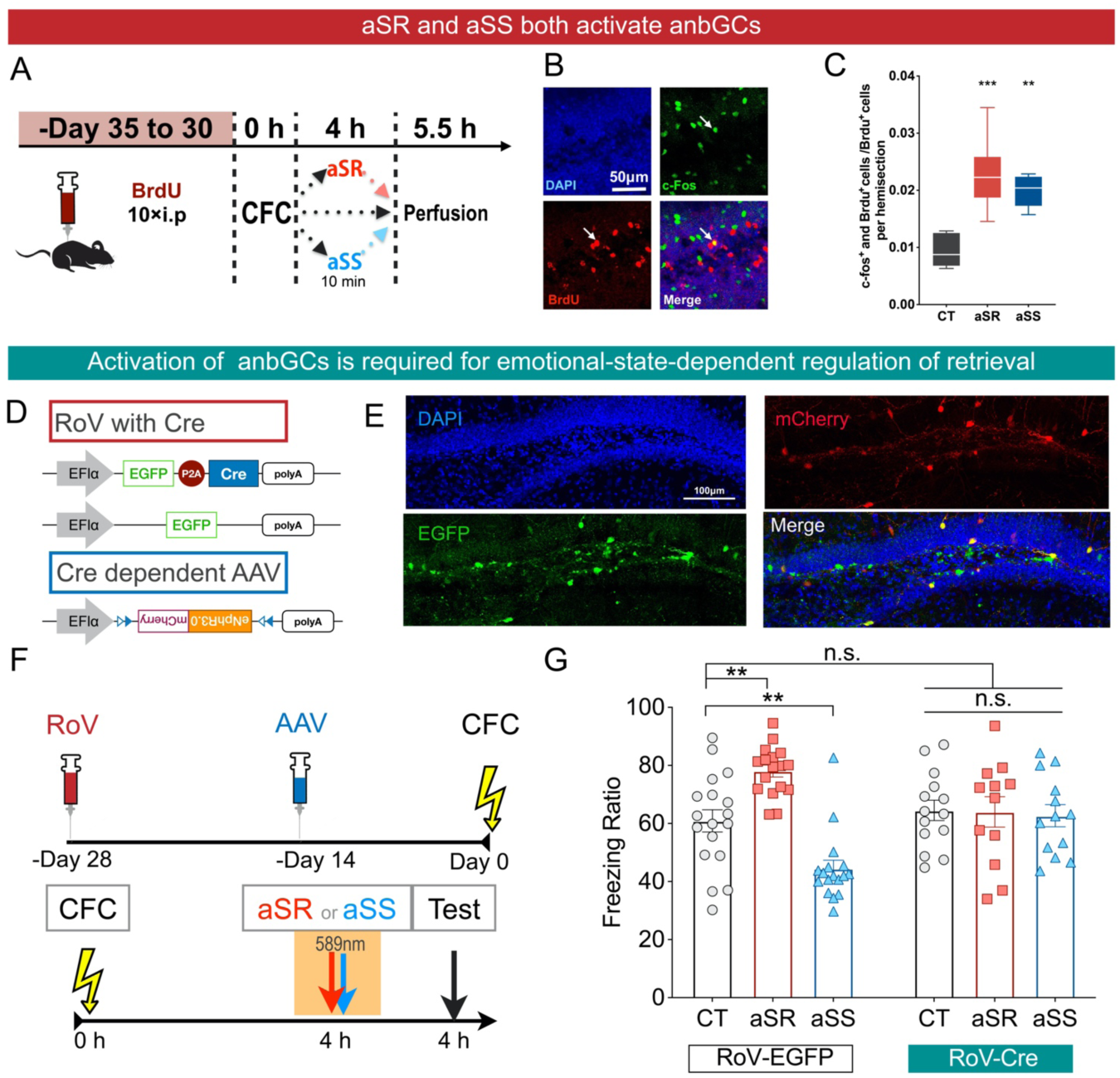
Acute social experiences regulate memory retrieval through activation of adult newborn granule cells (anbGCs). **(A)** Experimental schedule of retrieval-activated anbGC labelling. **(B)** Representative images of retrieval-activated anbGCs (labelled by c-Fos and BrdU). **(C)** Quantification of the ratio of retrieval-activated anbGCs (n_CT_=5, n_aSR_=6, n_aSS_=5). **(D)** Double virus-mediated anbGC labelling with eNpHR3.0. **(E)** Representative images of RoV (EGFP) and AAV (mCherry) expression. **(F)** Experimental schedule of inhibiting anbGCs during acute social experiences. **(G)** 4 h memory without anbGC inhibition during acute social experiences (RoV-EGFP: n_CT_=18, n_aSR_=17, n_aSS_=17) and with anbGC inhibition during acute social experiences (RoV-Cre: n_CT_=14, n_aSR_=12, n_aSS_=13). **P < 0.01, ***P < 0.001; one-way analysis of variance (ANOVA) with Dunnett’s test (C), or two-way analysis of variance (ANOVA) with Tukey’s test (G). Graphs show means ± SEM. See also Figures S5 and S6.

### anbGCs mediate the effects of aSR and aSS through bidirectional modulation of engram cell reactivation

Since reactivation of engram cells in the dDG is strongly correlated with memory retrieval performance (Kitamura et al., 2017; Reijmers et al., 2007; Roy et al., 2016), we assayed engram cell reactivation in memory retrieval with acute social experiences. Engram cells in the dDG were labelled with mCherry during the training sessions by utilizing the c-fos promoter-driven Tet-off system, as reported in previous studies (Kitamura et al., 2017; Liu et al., 2012; Reijmers et al., 2007), and the system was switched off by i.p. doxycycline administration 1.5 h before retrieval (Figure 4A). To visualize retrieval-activated neurons, we perfused mice 1.5 h after the recall test for c-Fos staining (Figures 4A and 4B; Figure S7). The ratio of retrieval-reactivated engram cells (mCherry and c-Fos co-labelled) to total engram cells (mCherry positive) was significantly increased with aSR and decreased with aSS (Figure 4C). Thus, aSR-dependent enhancement of memory corresponds to the reactivation of more engram cells upon retrieval, while aSS-dependent decrease of memory corresponds to fewer reactivated engram cells.

**Figure 4.**
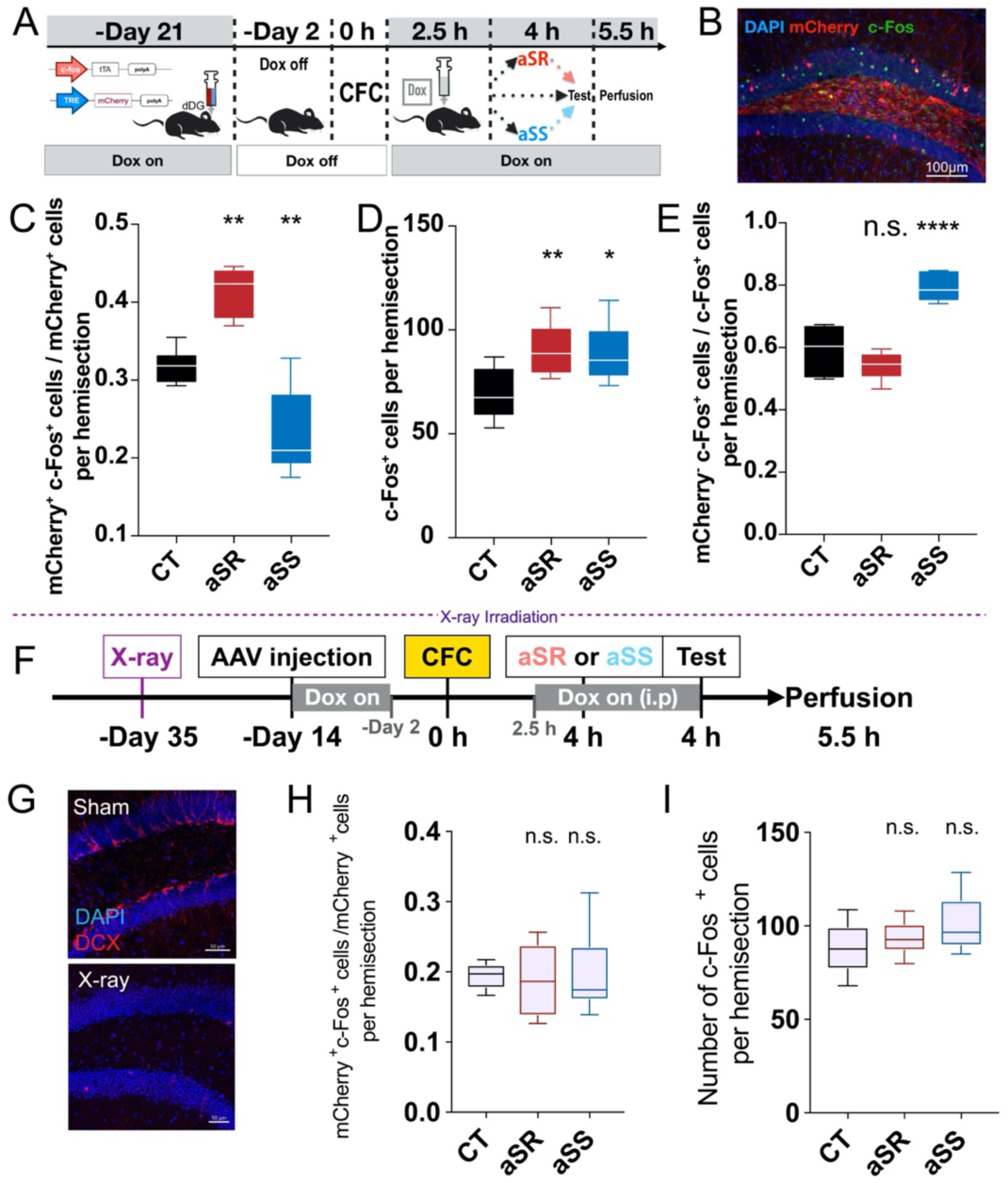
Acute social experiences modulate engram cell reactivation through anbGCs. **(A)** Experimental design for testing engram reactivation with acute social experiences. **(B)** Representative image of engram cells (mCherry labelled) and retrieval-activated cells (c-Fos labelled) in the dDG. **(C)** Quantification of the engram reactivation ratio (n_CT_=6, n_aSR_=6, n_aSS_=7). **(D)** Quantification of the retrieval-activated cells (n_CT_=6, n_aSR_=8, n_aSS_=6). **(E)** Quantification of the ratio of retrieval-activated non-engram cells (n_CT_=6, n_aSR_=6, n_aSS_=7). **(F)** Experimental design for testing engram reactivation with acute social experiences in mice with X-ray irradiation. **(G)** Representative images of anbGCs (DCX) in the dDG. **(H)** Quantification of the engram reactivation ratio in mice with X-ray irradiation (n_CT_=5, n_aSR_=6, n_aSS_=6). **(I)** Quantification of the retrieval-activated cells in mice with X-ray irradiation (n_CT_=5, n_aSR_=6, n_aSS_=6). *P < 0.05, **P < 0.01, ****P < 0.0001; one-way analysis of variance (ANOVA) with Dunnett’s test. Graphs show means ± SEM. See also Figure S7.

It is interesting to note that the total number of retrieval-activated c-Fos cells in the dDG was increased in both the aSR and aSS groups (Figure 4D). However, neurons activated by aSS had a higher ratio of non-engram cells (mCherry negative) compared with neurons activated by aSR (Figure 4E), suggesting that aSR enhanced the signal-to-noise ratio of memory by facilitating retrieval-dependent recruitment of engram cells, while aSS reduced the ratio by hindering recruitment and activating more non-engram cells during retrieval.

To investigate whether the observed modulation of engram cell reactivation depends on anbGCs, we examined the reactivation pattern of engram cells in mice irradiated with X-rays (Figures 4F and 4G). We found that blockade of neurogenesis could abolish acute social experience-mediated regulation of the engram cell reactivation pattern as well as the effect of increasing total dDG activity (Figures 4H and 4I).

### aSR and aSS activate distinct anbGC populations and trigger opposite modulation of dDG activity

Since the impacts of both aSR and aSS rely on activating anbGCs, the question arises as to how such activation leads to opposing effects. We examined whether different anbGC populations are involved. Using the double-virus system, we inhibited anbGCs during training so that they were excluded from encoding memory (Figure 5A). With such manipulation, we found that aSR failed to regulate memory retrieval (Figure 5B), while aSS remained effective (Figure 5C). This result is consistent with the idea of activating and modulating two distinct populations of anbGCs, with aSR acting through memory engram anbGCs while aSS activates non-engram anbGCs.

**Figure 5.**
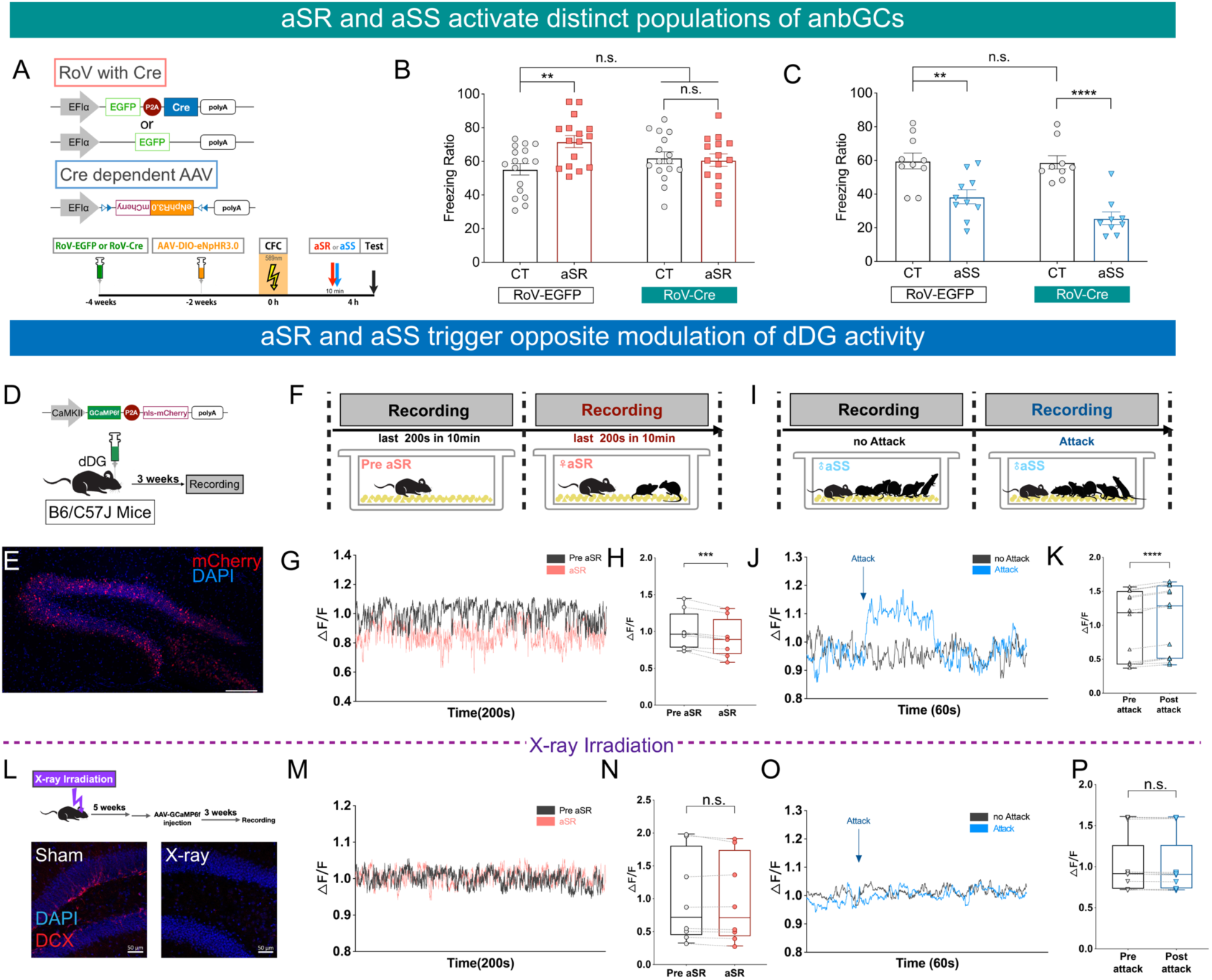
Acute social reward and stress activate distinct anbGC populations and trigger opposite modulation of dDG activity. **(A)** Experimental schedule of inhibiting anbGCs during CFC. **(B)** 4 h memory with aSR without anbGC inhibition during CFC (RoV-EGFP: n_CT_=17, n_aSR_=16) and with inhibition (RoV-Cre: n_CT_=16, n_aSR_=15). **(C)** 4 h memory with aSS without anbGCs inhibition during CFC (RoV-EGFP: n_CT_=10, n_aSS_=10) and with inhibition (RoV-Cre: n_CT_=9, n_aSS_=9). (**D)** Experimental schedule of calcium recording. **(E)** Representative images of AAV (mCherry) expression, scale bar: 100μm. **(F and I)** Experimental schedules of aSR (F) and aSS (I) during calcium recording. **(G and J)** Representative diagrams of calcium recording for aSR (G) and aSS (J). **(H)** Comparison of calcium signal between pre-aSR and aSR (n=8). **(K)** Comparison of calcium signal between pre-attack and post-attack (n=14). **(L)** Experimental schedule of calcium recording in mice with X-ray irradiation (left) and representative images of anbGCs (DCX) in the dDG (right). **(M and O)**, Representative diagrams of calcium recording for aSR (M) and aSS (O) in mice with X-ray irradiation. **(N)** Comparison of calcium signal between pre-aSR and aSR in mice with X-ray irradiation (n=8). **(P)** Comparison of calcium signal between pre-attack and post-attack in mice with X-ray irradiation (n=13). **P < 0.01, ***P < 0.001, ****P < 0.0001; two-way analysis of variance (ANOVA) with Tukey’s test (B and C), or paired two-tailed t test (others). Graphs show means ± SEM.

Given that anbGCs played an important role in regulating information processing in the dentate gyrus (DG) by regulating its overall activity (Anacker et al., 2018; Luna et al., 2019; Temprana et al., 2015), we hypothesized that aSR and aSS regulate memory by modulating dDG activity in different patterns. We measured dDG activity during acute social experiences by using AAV-expressed GCaMP6f driven by the CaMKIIα promoter to record calcium signals in the dDG with an implanted cannula implantation (Figures 5D and 5E). In the first trial, mice were allowed to move freely for 10 min unattended in a new cage with space for rearing. In the second trial, the mice were exposed for 10 min to either two female mice as aSR (Figure 5F) or five grouped male mice as aSS (Figure 5I). For aSR, by comparing the ratios (ΔF/F) of calcium signal in the last 200 s of these two trials, we found that dDG activity was significantly decreased after aSR (Figures 5G and 5H). For aSS, we quantified the calcium signals 60 s before and after the mouse was attacked (the average number of times the tested mouse was attacked during the second trial was 3.25, n=8) and found a sudden increase in dDG activity during each attack (Figures 5J and 5K).

We then investigated the relationship between this modulation of dDG activity and anbGCs. We found that when neurogenesis was blocked via X-ray irradiation (Figure 5L), both types of acute social experience failed to trigger changes in calcium activity (Figures 5M-5P). Collectively, these results showed that aSR and aSS have distinct, anbGC-dependent modulatory effects on dDG activity.

### dDG activity is sufficient to regulate and predict memory retrieval

The observed changes in dDG activity likely contribute to the regulation of memory retrieval. To further validate the roles of dDG activity in state-dependent memory plasticity, we altered dDG activity directly through optogenetic manipulation to mimic the regulatory effects of aSR and aSS as a way to confirm the relationship between dDG activity and sequential memory retrieval. Memory performance was significantly enhanced by inhibiting excitatory neuronal activity within the dDG through laser light stimulation of eNpHR3.0, driven by the CaMKIIα promoter, before retrieval (Figures 6A-6C). Conversely, an increase in dDG activity by stimulation of hChR2 reduced memory retrieval (Figures 6E-6G). As a control, manipulation of dDG activity did not influence the freezing performance of mice in a novel context (Figures 6D and 6H).

**Figure 6.**
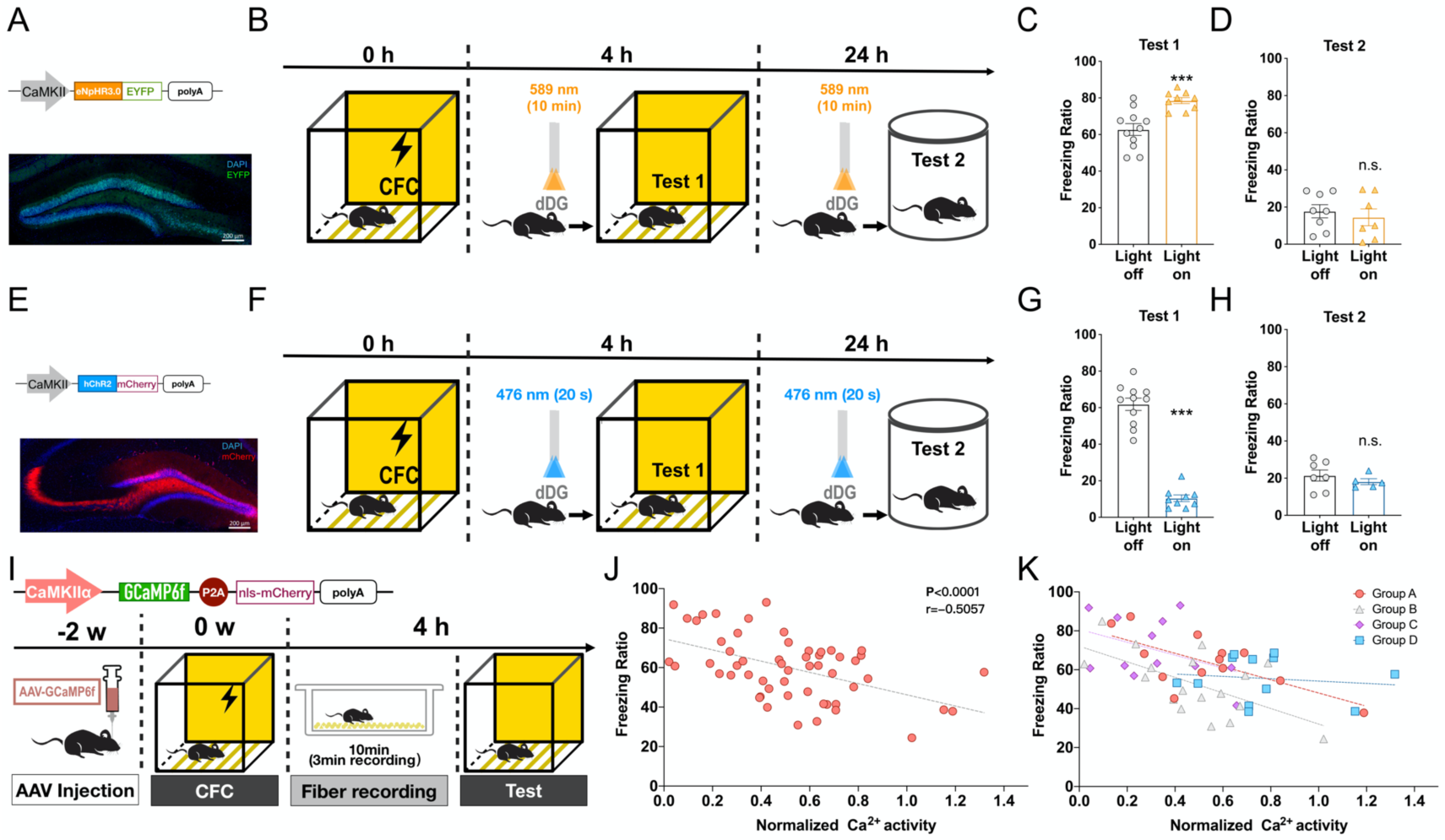
dDG activity is sufficient to regulate and predict memory retrieval performance. **(A and B)** Experimental schedule of optogenetic inhibition of dDG neurons before tests. **(C)** Freezing ratio in CFC context with optogenetic inhibition of dDG neurons before test (n_Light off_=11, n_Light on_=9). **(D)** Freezing ratio in novel context with optogenetic inhibition of dDG neurons before test (n_Light off_=8, n_Light on_=7). **(E and F)** Experimental schedule of optogenetic activation of dDG neurons before tests. **(G)** Freezing ratio in CFC context with optogenetic activation of dDG neurons before test (n_Light off_=11, n_Light on_=9). **(H)** Freezing ratio in novel context with optogenetic activation of dDG neurons before test (n_Light off_=7, n_Light on_=5). **(I)** Experimental schedule of calcium recording of the dDG before 4 h memory tests. **(J)** The correlation between calcium signal and freezing ratio in all mice (n=55), fit by linear regression model. **(K)** The separated data of (J) from four parallel groups (n_GroupA_=13, n_GroupB_=18, n_GroupC_=12, n_GroupD_=12).; ***P < 0.001; unpaired two-tailed t test (C, D, G and H). Graphs show means ± SEM.

To further investigate the relationship between overall dDG activity and retrieval efficacy, we tested the correlation between dDG neuronal activity prior to memory recall and retrieval performance. Four hours after training, Ca^2+^ activity in the dDG was recorded for 3 min in freely moving mice in their home cages with the expression of CaMKII-driven Gcamp6f. Immediately after dDG activity recording, mice were tested for memory in the training context (Figure 6I). By combining data from each individual mouse regarding dDG Ca^2+^ activity and memory performance, we found that dDG activity and memory retrieval efficacy had a significant negative correlation (Figures 6J and 6K), which was consistent with the results of dDG activity manipulation. Together, our data indicated that dDG activity was sufficient to both regulate and predict the efficacy of memory retrieval. These findings suggest that anbGC-triggered dDG activity modulation reflects the current emotional state and is a key process in the regulation of retrieval efficacy.

### Emotional state-dependent memory plasticity is biased in stress models and can be recused by promoted neurogenesis

Given that depression is a typical emotional disorder and that hippocampal neurogenesis is impaired in many depression models (Miller and Hen, 2015; Santarelli and Hen, 2003), we next tested whether this emotional-state-dependent memory plasticity is affected in stress models for the study of depression-related phenotypes.

We employed two procedures to induce depression-like behaviors in mice, lipopolysaccharide (LPS) injection (Figure 7A) and chronic forced swimming (CFS) (Figure 7G). Both treatments were associated with impairment in the open field test (Figures 7B and 7H) and in the tail suspension test (Figures 7C and 7I), two parameters for the establishment of depression phenotypes. In these two models, mice with depression-like behaviors showed negatively biased plasticity of memory retrieval; mice did not show enhanced memory retrieval after aSR, while the effect of aSS remained intact (Figures 7D and 7J). In support of this observation, aSR-dependent downregulation of dDG activity was also blocked (Figures 7E, 7F, 7K, 7L). Furthermore, this biased regulation did not result from alterations in social interaction time during the social reward period (Figure S9). These data are consistent with the biased memory theory in depression patients (Dillon and Pizzagalli, 2018; Disner et al., 2011; Mathews and MacLeod, 2005), namely, that those patients have improved negative memory but impaired positive memory.

**Figure 7.**
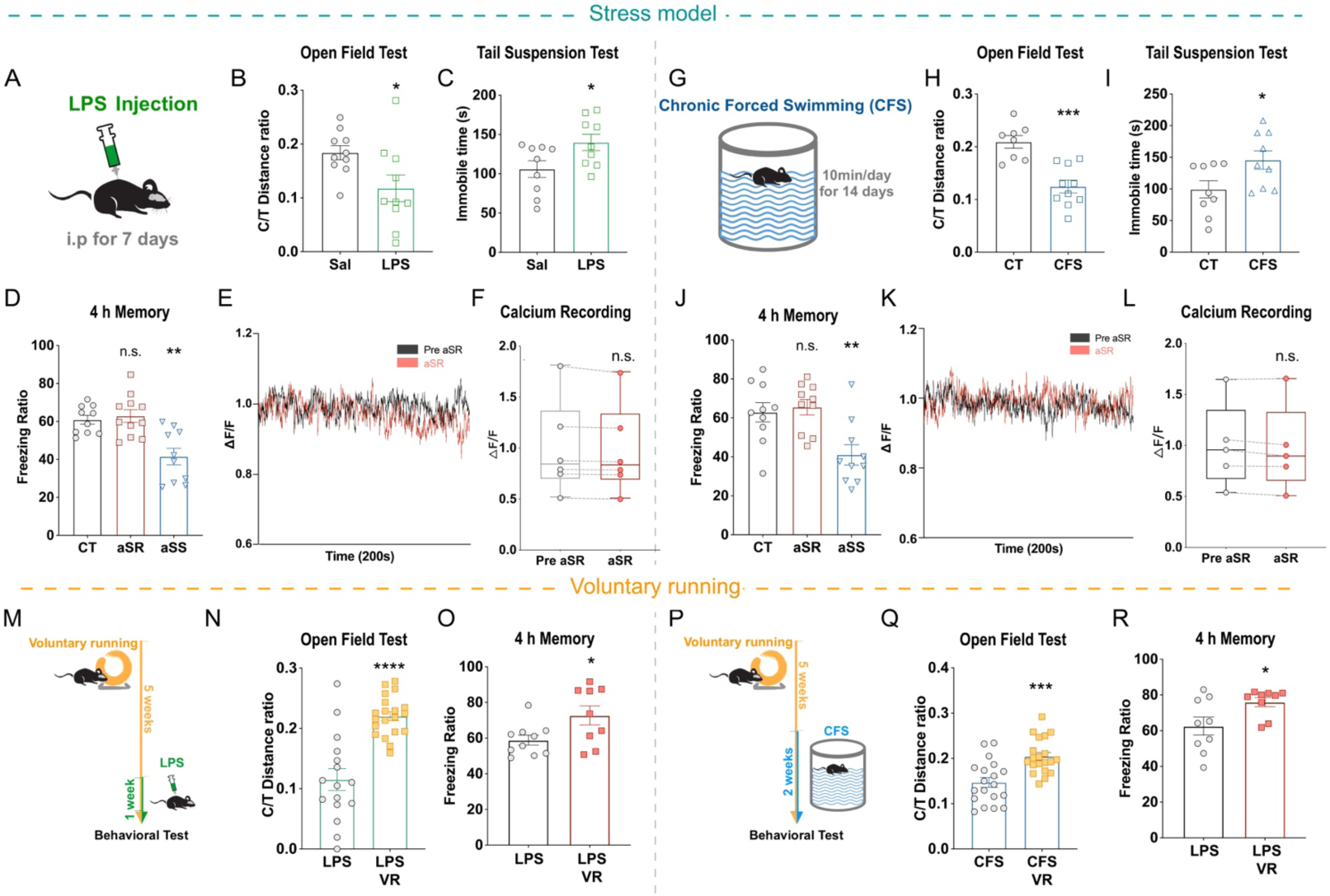
anbGC-mediated regulation of retrieval is biased in stress models, and the bias can be ameliorated by increasing neurogenesis. **(A)** Stress model induced in mice by LPS injection. **(B)** Open field tests in LPS-injected mice (n_Sal_=10, n_LPS_=10). **(C)** Tail suspension tests in LPS-injected mice (n_Sal_=9, n_LPS_=9). **(D)** 4 h memory with acute social experiences in LPS-injected mice (n_CT_=10, n_aSR_=11, n_aSS_=10). **(E)** Representative diagram of calcium recording for aSR in LPS-injected mice. **(F)** Comparison of calcium signal between pre-aSR and aSR in LPS-injected mice (n=6). **(G)** Stress models induced in mice by chronic forced swimming (CFS). **(H)** Open field tests in CFS mice (n_CT_=8, n_CFS_=10). **(I)** Tail suspension tests in CFS mice (n_CT_=9, n_CFS_=9). **(J)** 4 h memory with acute social experiences in CFS mice (n_CT_=10, n_aSR_=10, n_aSS_=10). **(K)** Representative diagrams of calcium recording for aSR in CFS mice. **(L)** Comparison of calcium signal between pre-aSR and aSR in CFS mice (n=5). **(M and P)** Experimental schedules of inducing stress models in mice with voluntary running (VR). **(N)** Open field tests in LPS-injected mice with VR (n_LPS_=16, n_LPS-VR_=20). **(O)** Regulation of aSR on 4 h memory in LPS-injected mice with VR (n_CT_=10, n_aSR_=9). **(Q)** Open field test in CFS mice with VR (n_CFS_=19, n_CFS-VR_=20). **(R)** Regulation of aSR on 4 h memory in CFS mice with VR (n_CT_=9, n_aSR_=9). *P < 0.05, **P < 0.01, ***P < 0.001, ****P < 0.0001; one-way ANOVA with Dunnett’s test (D and J), paired two-tailed t test (F and L), or unpaired two-tailed t test (others). Graphs show means ± SEM. See also Figures S9 and S10.

Given that activating anbGCs or increasing neurogenesis can ameliorate depression-like behaviors in rodents(Hill et al., 2015; Snyder et al., 2011; Tunc-Ozcan et al., 2019) and that hippocampal neurogenesis is required for antidepressants to be effective in treating depression(Miller and Hen, 2015; Santarelli and Hen, 2003), we next evaluated whether the promotion of neurogenesis could help mice retain the ability to use positive stimuli to regulate memory retrieval after LPS injection or CFS. We found that five weeks of voluntary running could effectively promote neurogenesis (Figures S10A-S10C). Indeed, after voluntary running, mice subjected to LPS injection or CFS showed ameliorated performance in open field tests (Figures 7N and 7Q) and the enhancement of memory retrieval after aSR remains effective (Figures 7M, 7O, 7P, 7R). Our data suggested that in mice with depression-like behaviors, biased acute social experiences-based plasticity of retrieval can be ameliorated by increasing neurogenesis.

## Discussion

Determining how the hippocampus integrates different emotional information in memory processing is crucial for our understanding of how emotion affects animal behavior. Previous studies have revealed that acute emotional experiences can affect memory through hormone- or catecholamine-triggered modification of synaptic strength (Fernández and Morris, 2018; Hu et al., 2007; Kim and Diamond, 2002; De Quervain et al., 1998; Tak et al., 2007; Takeuchi et al., 2016). However, how the hippocampus integrates diverse experiences and how such modification is reflected in specific memory engrams remain elusive. In the current work, we sought to determine how emotional states triggered by acute social experiences affect memory and to identify the neural basis of this emotional state-dependent memory plasticity.

First, we found that retrieval of a hippocampal memory could be up- or downregulated significantly by acute social events prior to it within a short time window of 30 min, that aSR led to an enhanced while aSS led to an attenuated memory. Such effects were confined to retrieval and did not affect memory acquisition (see Figures 1A-1F). The pervasive nature of such subtle daily social experiences and their impact on memory retrieval clearly demonstrate that memory retains plastic even after consolidation. Such plasticity allows the efficacy of memory expression to fluctuate from moment to moment depending on the state immediately before the recall (see Figures 1G-1K), thereby helping animals to achieve optimal behavioral outcomes when facing diverse situations.

Second, flexible retrieval of memory is mediated by the activation of two distinct populations of anbGCs: aSR activates training-activated anbGCs, leading to the recruitment of more engram cells in retrieval, while aSS activates non-training-activated anbGCs, leading to the recruitment of fewer engram cells. The involvement of anbGCs is demonstrated by two observations. First, we observed that aSR and aSS are both capable of activating anbGCs. Second, we found that retrieval plasticity was abolished by the blockade of neurogenesis (see Figure 2) or optogenetic inhibition of anbGC activity (see Figures 3D-3G). The involvement of two distinct populations of anbGCs is supported by the following three observations: (1) Excluding anbGCs from engram cells impedes only the effect of aSR and leaves the effect of aSS intact (see Figures 5A-5C); (2) aSR and aSS exhibited opposite modulatory effects on dDG neuronal activity, which is mediated by anbGCs, and such modulation is sufficient to regulate and predict performance in memory retrieval (see Figure 5 and Figure 6); (3) In stress models, only aSR-induced enhancement in retrieval was impaired while the aSS effect remains intact, and such impairment could be ameliorated through voluntary running-promoted neurogenesis (see Figure 7). These three lines of evidence suggest that anbGCs confer plasticity of memory retrieval through two distinct neuronal mechanisms that are activated by positive and negative states. This notion is consistent with recent reports that the activity of anbGCs plays an important role in regulating information processing in the dentate gyrus through distinct signaling pathways by either activating GABAergic neurons to suppress activity (Temprana et al., 2015) or regulating excitatory synapses via anbGC-activated glutamate receptors (Luna et al., 2019).

Taken together, our data demonstrate that the acute emotional state of animals can bidirectionally regulate memory retrieval by activating distinct populations of anbGCs. These data suggest that activation of distinct populations of anbGCs leads to bidirectional modulation of dDG activity that, in turn, affects the reactivation efficacy of engram cells. Such changes in reactivation efficacy determine memory retrieval plasticity. Thus, anbGCs represent a convergence point in the transformation of both positive and negative acute emotional events into appropriate levels of memory retrieval to adapt to an ever-changing environment.

## Acknowledgments

We thank all the members of the Zhong lab for their support. This work was supported by grants from the National Science Foundation of China (91332207 and 91632301, to Y.Z.), the National Basic Research Project (973 program) of the Ministry of Science and Technology of China (2013cb835100, to Y.Z.), the Beijing Municipal Science and Technology Commission (Z161100002616010, to Y.Z.), and the Tsinghua-Peking Joint Center for Life Sciences.

## Author Contributions

B.L. and Y.Z. conceived and designed the study and wrote the paper. B.L. and B.K. performed most of the experiments. B.L. and H.C. performed behavioral experiments and analyzed the data. W.L. performed calcium recording. B.K. and W.L. conducted surgeries. B.K., Y.H. and B.L. performed immunohistochemistry and cell counting. J.M. and S.S. conducted in vitro electrophysiological experiments.

## STAR Methods

### RESOURCE AVAILABILITY

#### Lead Contact

Further information and requests for resources and reagents should be directed to and will be fulfilled by the Lead Contact, Yi Zhong.

#### Materials Availability

This study did not generate new unique reagents.

#### Data and Code Availability

No code is generated in this study. Data generated in this study are available upon reasonable requests.

### EXPERIMENTAL MODEL AND SUBJECT DETAILS

#### Animals

All experimental procedures were approved by Institutional Animal Care and Use Committee (IACUC) of Tsinghua University, and were performed using the principles outlined in the Guide for the Care and Use of Laboratory Animals of Tsinghua University. All mice were purchased and maintained under standard conditions by the Animal Research Center of Tsinghua University. The male mice used throughout the whole study were 8-12 weeks old. The female mice used in social reward treatments were 4-8 weeks old. The juvenile mice used in social reward treatments were 4 weeks old. The retired male CD-1 breeder mice used in social stress treatments were 6-10 months old.

### METHOD DETAILS

#### Viral constructs

The AAV_9_-c-Fos-tTA-pA, AAV_9_-TRE-mCherry, pAAV-EF1α-DIO-caspase3-P2A-GFP, AAV_9_-EF1α-DIO-eNpHR3.0-EYFP, AAV_9_-EF1α-DIO-eNpHR3.0-mCherry, AAV_9_-EF1α-DIO-mCherry, AAV_9_-CaMKIIα-eNpHR3.0-EYFP, AAV_9_-CaMKIIα-hChR2(H134R)-mCherry and AAV_9_-CaMkIIα-GCaMP6f-P2A-nls-dTomato viruses were acquired from Vigene. The pRoV-EF1α-EGFP-2A-Cre and pRoV-EF1α-EGFP viruses were acquired from OBiO. The viral concentration was 1.42 × 10^13^ viral genomes (vg) ml^−1^ for AAV_9_-c-Fos-tTA-pA, 1.02 × 10^13^ vg ml^−1^ for AAV_9_-TRE-mCherry, 6.59 × 10^13^ vg ml^−1^ for pAAV-EF1α-DIO-caspase3-P2A-GFP, 2.18 × 10^13^ vg ml^−1^ for AAV_9_-EF1α-DIO-eNpHR3.0-EYFP, 5.29 × 10^13^ vg ml^−1^ for AAV_9_-EF1α-DIO-eNpHR3.0-mCherry, 1.55 × 10^14^ vg ml^−1^ for AAV_9_-EF1α-DIO-mCherry, 1.85 × 10^13^ vg ml^−1^ for AAV_9_-CaMKIIα-eNpHR3.0-EYFP, 9.52 × 10^13^ vg ml^−1^ for AAV_9_-CaMKIIα-hChR2(H134R)-mCherry, 7.24 × 10^13^ vg ml^−1^ for AAV_9_-CaMkIIα-GCaMP6f-P2A-nls-dTomato, 5.13 × 10^7^ vg ml^−1^ for pRoV-EF1α-EGFP-2A-Cre, and 1.21 × 10^9^ vg ml^−1^ for pRoV-EF1α-EGFP. Viruses were subdivided into aliquots and stored at −80°C until use.

#### Stereotactic injection and optical fiber implant

Mice were anaesthetised with 0.5% sodium pentobarbital (70mg/kg, 350ul/25g). Bilateral craniotomies were performed using a 0.5 mm diameter drill, and 400 nl of the virus was injected into the dDG using a 10 μl nanofil syringe controlled by UMP3 and Micro4 system (WPI) with a speed of 60 nl/min. The dDG injections were bilaterally targeted to −2.2 mm AP, ± 1.3 mm ML, and −2.0 mm DV. After the injection, the needle remained in place for 10 min to ensure that the virus spread to the targeted area before it was slowly withdrawn. For mice need fiber implanting, after withdrawing the needle, a Doric fiber-optic patch cord (200 μm core diameter; Doric Lenses) was precisely cut to the optimal length and lowered above the injection site (–2.2 mm AP; ± 1.3 mm ML; –1.8 mm DV). A jewelry screw was screwed into the skull to provide an extra anchor point. Dental cement (Teets Cold Cure; A-M Systems) was applied to secure the optical fiber implant. After surgery, the mice were allowed to recover for at least 2 weeks before all subsequent experiments were performed

#### Drugs

##### BrdU

BrdU (Sigma) was dissolved in 0.9% saline containing 0.007 mol/L NaOH and injected (100 mg/kg, i.p.) into mice twice per day at 12 h intervals for 5 consecutive days.

##### Temozolomide (TMZ)

TMZ (Sigma) was dissolved in 0.9% saline containing 10% DMSO and injected (25 mg/kg, i.p.) per day for 4 consecutive days. A total of four rounds of treatment (one week apart) were administered to mice (Garthe et al., 2009).

#### Voluntary running

Mice in the voluntary running group were given voluntary access to two running wheels (13 cm diameter) placed in their home cages. Mice in the sedentary group were similarly housed but were not given running wheels (Akers et al., 2014; Van Praag et al., 1999).

#### Local X-ray irradiation

Mice were anaesthetised with Avertin (250 mg/kg, i.p.) and before being placed under the X-ray irradiation apparatus (Radsource). A lead shield was used to cover the animal’s entire body except the head during irradiation. The dose rate was approximately 0.86 Gy/min at a source-to-skin distance of 13 cm. The irradiation lasted 700 s and delivering a total of 10 Gy (Kitamura et al., 2009).

#### Acute social reward/stress treatment

For social reward treatment used throughout the whole study, we placed two female mice, aged between 4 and 8 weeks, in a cage (W, 16 cm; D, 28 cm; H, 13 cm) with soft packing beforehand for 5-10 min to acclimate the mice to the new environment. Then, we placed the male mouse to be tested in contextual fear conditioning into the cage for 10 min. For social stress treatment used throughout the whole study, we placed the male mouse to be tested into a home cage containing five male mice for 10 min. For social reward treatment using juvenile, we placed two juvenile mice, aged 4 weeks, in the cage (W, 16 cm; D, 28 cm; H, 13 cm) with soft packing beforehand for 5-10 min to acclimate the mice to the new environment. Then, we placed the male mouse to be tested into the cage for 10 min. For social stress induced by social defeat, we placed the male mouse to be tested into a home cage containing a retired male CD-1 breeder mouse (6-10 months old) for 10 min.

#### Behavioral testing

##### Social interaction test

Mice were subjected to the social reward procedure described above. Experimenters monitored the whole process and recorded the time of intimate interactions between the male mouse and the female mice. Intimate interaction was defined as actions in which the male mice sniffs, chases, or attempts to mate with the female. The experimenters were blinded to the groups during the tests.

##### Contextual Fear Conditioning (CFC)

Fear conditioning was performed on male mice. For fear conditioning, the context was a chamber (W, 18 cm; D, 18 cm; H, 29 cm) and a grid floor that consisted of 16 stainless steel rods. Subject mice were placed in the conditioning chamber for 2 min before being given 5 foot shocks (2 s, 0.8mA, 1 min apart). 4 h after training, mice were placed into the original context and were allowed to explore for 180 s in order to monitor their freezing behavior. The same procedure was conducted at 24 h, 48 h or 7 days after training respectively. For Figures. 1J and 1K, we set two sequential retrieval tests at 4 h after training and 6 h after training. 4 h after training, the mice’s memory performance was first tested for 120 s in the chamber after a 10 min social experience. 2 h later, mice were tested again for 180s after another 10 min social experience.

##### Open Field Test

The open field test was conducted in an open plastic arena (50 cm × 50 cm × 40 cm). Mice were first placed in the peripheral area with their head towards the wall. Exploration time during 10 min in the peripheral and central areas, respectively, was measured using an automated video-tracking system (Anymaze software). The central field is defined as a 20 cm × 20 cm area in the middle of the arena.

##### Tail Suspension Test

Mice were suspended by their tails with scotch tape, in a position avoiding them escaping or holding on to nearby surfaces. The immobile time was recorded manually during the 5 min test. The experimenters were blinded to the groups during the tests.

#### Laser Delivery

##### Inhibition of anbGCs during social experiences

A 589 nm laser was bilaterally delivered by two optic fibers for optogenetical inhibition. Mice to be tested in CFC were first anaesthetised by isoflurane for fiber implantation. Then, they experienced either social reward or stress for 10 min. The timing started when the mice recovered from the anaesthesia and acted normally. After a 10 min test, mice were anaesthetised again to unplug the fiber. After mice recovered from anaesthesia for 3-5 min, they were tested for memory performance. Control mice were not infected with eNpHR3.0 but were still implanted with the cannula delivering light into the dorsal hippocampus.

##### Inhibition of anbGCs during training

Mice were anaesthetised before and after light delivery. Light from the 589 nm laser was bilaterally delivered to inhibit anbGCs throughout the training procedure.

##### Inhibition or activation of DG neurons

Mice were anaesthetised before and after light delivery. Mice were placed into a box that was distinguishable from the training context. Light from a 589 nm laser was bilaterally delivered for inhibition, while hChR2 was stimulated bilaterally at 12 Hz (10 ms pulse width) using a 473 nm laser (10–15 mW). hChR2 stimulation was conducted for only 20-30 s during the 10 min assessment.

#### Calcium Recording

Calcium signals were measured three weeks after viral infection with AAV_9_-CaMkIIα-GCaMP6f-P2A-nls-dTomato and implantation of a single lateral optical fiber at the dDG. A 40-60 s baseline recording of the environment was taken, with the light off, prior to recording each conditioned mouse. Mice to be recorded were first anaesthetised using isoflurane for fiber implantation. When testing the change of dDG activity caused by social interaction, mice were allowed to move freely, unattended, in a non-homecage box for 10 min, and the final 200 s signal was recorded. This procedure was followed by another 10 min recorded exposure of the mouse being recorded to either a pair of female mice (social reward treatment) or five male mice (social stress treatment). Social reward treatment was recorded during the last 200 s of the total 10 min trial, while in the social stress treatment the no-attack period and the attack periods (before and after the attack) were recorded during the 10 min trial. When testing dDG activity to predict behavior performance, mice were allowed to move freely, unattended, in a non-homecage box, and their calcium signal during the whole 3 min was recorded.

Data of the calcium recording were presented as ΔF/F versus time (s). ΔF/F was computed as (F(t) – F_0_)/F_0_, where F(t) was the raw calcium signal and F_0_ was the mean baseline fluorescence of the environment during the 40-60 s preceding each recording. The last 200 s of social reward treatment were compared with the last 200 s of the unattended state. For the social stress treatment, attack periods were compared with no-attack periods.

#### Electrophysiology and Immunohistochemistry for recorded slices

Double-virus system was used to drive the expression of the optogenetic tool, eNpHR3.0 that targets age-specific anbGCs in conjunction with pRoV-EF1α-EGFP-2A-Cre, injected into the dDG at 4 weeks, and AAV_9_-EF1α-DIO-eNpHR3.0-EYFP, at 2 weeks, before in vitro electrophysiological recording. Brains were removed and acute cortical slices (300–350 μm) were prepared using a vibratome (VT1200S, Leica) in chilled choline solution containing (in mM): 120 choline chloride, 2.6 KCl, 26 NaHCO_3_, 1.25 NaH_2_PO_4_, 7 MgSO_4_, 0.5 CaCl_2_, 1.3 ascorbic acid, 15 D-glucose, bubbled with 95% O_2_ and 5% CO_2_ on. Slices were kept in artificial cerebral spinal fluid containing (in mM): 126 NaCl, 3 KCl, 26 NaHCO_3_, 1.2 NaH_2_PO_4_, 1.3 MgSO_4_, 2.4 CaCl_2_, 10 D-glucose, bubbled with 95% O_2_ and 5% CO_2_, recovered at 32 °C for 1 hr and then stored at room temperature before recording. All slice recordings were performed at 34 °C. An upright microscope (Olympus) equipped with epifluorescence and infrared DIC illumination, a charge coupled device camera and two water immersion lenses (x10 and x60) was used to visualize and target recording electrodes to eGFP-expressing neurons located in dentate gyrus. Short pulses of light (590 nm, 700mA) were generated by a digital micro-mirror device (Polygon400, Mightex) integrated into the optical path of the microscope through the 10X objective (Figure S5). PolyScan2 software (Mightex) was used to control the location and shape of the area of stimulation. Microelectrodes (5–8MΩ) were filled with an intracellular solution containing (in mM): 110 potassium-gluconate, 20 KCl, 2 MgCl_2_, 0.2 EGTA, 10 HEPES, 4 MgATP, 0.4 Na_3_GTP and 0.3% neurobiotin (Invitrogen) (pH 7.25 and 295 mOsm). Whole-cell recordings were performed with an Axon Multiclamp 700B amplifier and pCLAMP10.6 (Molecular Devices). Series resistance was 10–30 MΩ and periodically monitored. Data were analyzed using Clampfit.

After recording, slices were incubated in 4% paraformaldehyde in PBS at 4 °C overnight, and then stained with primary antibody chicken anti-GFP (Aves, 1:1000) at 4 °C for 2 days. Slices were incubated with secondary antibody, including Alexa 488 donkey anti-chicken (Jackson Lab, 1:500) and 546–conjugated streptavidin (Invitrogen) at 4 °C overnight. Z-series images were acquired using a confocal laser scanning microscope (Olympus FV3000) (Figure S5).

#### Engram labeling

In order to label the engram cells formed in the dDG during training, we injected a mixture of AAV_9_-c-Fos-tTA-pA and AAV_9_-TRE-mCherry into mice (Liu et al., 2012)(Kitamura et al., 2017). Following virus injection, the mice were fed with diet containing 40 mg/kg doxycycline (Dox). To label the engram cells formed during the CFC, the mice were taken off Dox for two days. The mice were then subjected to CFC, and injected with Dox (i.p., 5mg/ml, 10μl/g mouse) 1.5 h before memory retrieval. When Dox treatment was not stopped in mice, no detectable labeled cells can be seen in the dDG (Figure S7).

#### Histology

##### Tissue Preparation

1.5 h after acute social experiences, retrieval or training, mice were deeply anaesthetised with 2% pentobarbital (160mg/kg,10ul/g) and transcardially perfused, first with cold PBS and then with cold 4% paraformaldehyde (PFA). The brains were extracted, postfixed in PFA overnight at 4°C, transferred to a 30% sucrose solution, and then stored at 4°C for 3 days. Next, the brains were coronally sectioned at 60-μm through the dentate gyrus using a freezing microtome.

##### Immunohistochemistry

Floating sections were used in all the following immunostaining experiments. Unless otherwise stated, all incubations occurred at room temperature.

For c-Fos and DCX immunostaining, sections were washed 3 times in 1× PBS and rinsed in 1% Triton X-100 for 15 min before a blocking step in PBS with 0.5% Triton X-100 and 10% normal donkey serum for 1 hr. Incubation with primary antibody was performed at 4°C for 48 hr (rabbit anti-doublecortin, Cell Signaling Technology, 1:500; guinea pig anti-c-Fos, Synaptic Systems, 1:500) in PBS with 0.5% Triton X-100 and 1% normal donkey serum. Sections were then washed 3 times in PBS and incubated with secondary antibody (donkey anti-rabbit IgG Alexa Fluor 647, donkey anti-GP IgG Cy3, Jackson ImmunoResearch, 1:500) for 2 hr. Sections were then washed 3 times in PBS and incubated in DAPI that was diluted in PBS (1:3000) for 10 min. Next, sections were again washed in PBS for 3 times before being mounted onto slides and coverslipped with antifade mounting medium (Invitrogen).

For BrdU immunochemistry, slices were washed 3 times in PBS and then incubated in 2M HCl for 30 min at 37°C. Next, HCl was removed, slices were washed 3 times in PBS and then neutralised with 0.1 M sodium borate buffer for 10 min. Sodium borate buffer was removed and slices were washed 3 times in PBS. The following steps were identical to those described for the c-Fos/DCX immunostaining, as described above. The sections were stained with primary antibody (rat anti-BrdU, abcam, 1:500) and secondary antibody (donkey anti-rat IgG Alexa Fluor 647, Jackson ImmunoResearch, 1:500).

##### Imaging and Quantification

Fluorescence was detected using a Zeiss LSM 880 with Airyscan for Figure 2F and Figure 3E, and an LSM 710 META was used for imaging the other figures with the aid of the ZEN software (black edition). Images were acquired with a 20× objective and colocalization was confirmed by a 3-D reconstruction of z series images.

For cell counting, six unilateral dDG regions across all the sections from one brain sample were chosen randomly. Counting was performed manually using the ZEN software (blue edition). Reported cell counts were calculated by averaging the results from all six sections.

#### Induction of Stress Models

##### Chronic Forced Swimming-Induced Stress Model

Mice were individually placed in an open cylindrical container (diameter 20 cm, height 30 cm), containing 20 cm depth of room temperature water. Each mouse was forced to swim for 10 min each day for 14 consecutive days. After swimming, mice were dried in a separate box and were then placed back to their home cage (Porsolt et al., 1977).

##### LPS-Induced Stress Model

Lipopolysaccharide (LPS) (Sigma) was dissolved in 0.9% saline and injected (0.5 mg/kg, i.p.) into mice every day for 7 consecutive days(Adzic et al., 2015).

### QUANTIFICATION AND STATISTICAL ANALYSIS

All experiments of behavioral tests and imaging were performed and analyzed blindly to the experimental condition. Statistical tests were performed using GraphPad Prism 7.0. No statistical methods were used to predetermine sample sizes, but our sample sizes are similar to those reported in previous publications. Student’s t test was used for the comparison between two groups. One-way ANOVA with Dunnett’s test was used for comparing a number of groups with a single control group. Two-way ANOVA with Tukey’s test was used for comparison of multiple groups with two factors. All tests were two-tailed. The sample size and statistical tests used are indicated in the figure legends. The sample size “n” represents number of mice. A full report of P values and statistics were summarized in **Statistics Information.** A P value less than 0.05 was considered statistically significant. The data are shown as the mean ± standard error of the mean (SEM). *p < 0.05; **p < 0.01; ***p < 0.001; ****p < 0.0001; n.s., nonsignificant (p > 0.05).

### KEY RESOURCES TABLE

| REAGENT or RESOURCE | SOURCE | IDENTIFIER |
| --- | --- | --- |
| Antibodies |  |  |
| primary antibody chicken anti-GFP | Aves Labs | GFP-1010<br>RRID:<br>AB_2307313 |
| Rabbit anti-doublecortin | Cell Signaling Technology | 4604<br>RRID:AB_561007 |
| Guinea pig anti-c-fos | Synaptic Systems | 226 005,<br>RRID:AB_2800522 |
| donkey anti-rabbit IgG Alexa Fluor 647 | Jackson ImmunoResearch | 711-605-152,<br>RRID:AB_2492288 |
| donkey anti-Guinea Pig IgG Cy3 | Jackson ImmunoResearch | 706-165-148,<br>RRID:AB_2340460 |
| donkey anti-rat IgG Alexa Fluor 647 | Jackson ImmunoResearch | 712-605-153<br>RRID:<br>AB_2340694 |
| Alexa 488 donkey anti-chicken | Jackson ImmunoResearch | 703-545-155<br>RRID:<br>AB_2340375 |
| Rat anti-BrdU | abcam | ab6326,<br>RRID:AB_30542<br>6 |
| **Bacterial and Virus Strains** |  |  |
| AAV9-c-Fos-tTA-pA | Vigene | N/A |
| AAV9-TRE-mCherry | Vigene | N/A |
| pAAV-EF1α-DIO-caspase3-P2A-GFP | Vigene | N/A |
| AAV9-EF1α-DIO-eNpHR3.0-EYFP | Vigene | N/A |
| AAV9-EF1α-DIO-eNpHR3.0-mCherry | Vigene | N/A |
| AAV9-EF1α-DIO-mCherry | Vigene | N/A |
| AAV9-CaMKIIα-eNpHR3.0-EYFP | Vigene | N/A |
| AAV9-CaMKIIα-hChR2(H134R)-mCherry | Vigene | N/A |
| AAV9-CaMKIIα-GCaMP6f-P2A-nls-dTomato | Vigene | N/A |
| pRoV-EF1α-EGFP-2A-Cre | OBiO | N/A |
| pRoV-EF1α-EGFP | OBiO | N/A |
| **Chemicals, Peptides, and Recombinant Proteins** |  |  |
| DAPI | Sigma | D9542 |
| Lipopolysaccharides | Sigma | L2880 |
| Sodium Chloride | Sigma | S5886 |
| Temozolomide | Sigma | PHR1437 |
| Paraformaldehyde | Sigma | 441244 |
| 5-Bromo-2'-deoxyunidine | Sigma | B5002 |
| Phosphate Buffer Saline | Solarbio | P1010 |
| Triton X-100 | Amresco | 0694 |
| ProLong Gold antifade reagent | Invitrogen | P36934 |
| 546-conjugated streptavidin | Invitrogen | S11225 |
| Hydrochloric Acid | Beijing Chemical Works | A0300003 |
| Sucrose | J&K Scientific | 971624 |
| Normal Donkey Serum | Jackson Immuno Research | 017-000-121 |
| Tissue Freezing Medium | Leica | 14020108926 |
| Sodium Tetraborate, Decahydrate | Beijing Biodee Biotechnology | S9640 |
| Ethanol | Beijing Tong Guang Fine Chemicals Company | 12404 |
| Pentobarbital | Animal Research<br>Center of<br>Tsinghua<br>University | N/A |
| Avertin | Animal Research<br>Center of<br>Tsinghua<br>University | N/A |
| Isoflurane | RWD Life Science | 217170702 |
| Experimental Models: Organisms/Strains |  |  |
| C57BL/6J | Jackson Labs | N/A |
| CD-1 | Charles River | N/A |
| Software and Algorithms |  |  |
| GraphPad Prism 7 | GraphPad<br>Software | http://www.graphpad.com/scientific-software/prism |
| Adobe Photoshop | Adobe.com | N/A |
| PolyScan2 software | Mightex | mightexbio.com |
| ANY-maze software | Stoelting Co. | anymaze.co.uk |
| FreezeFrame software | Actimetrics | N/A |
| ZEISS ZEN | Zeiss.com | N/A |
| Other |  |  |
| Doxycycline mouse food, custom | HFK Bio | N/A |
| Optic fiber implants, custom | Newdoon | N/A |
| Lasers for optogenetics | Changchun New<br>Industries<br>Optoelectronics<br>Technology | N/A |

#### Statistics Information

**Figure 1:**

**(B)** F (2, 39) = 16.29, P < 0.0001. CT vs. aSR (P = 0.0073), CT vs. aSS (P = 0.0232). **(C)** F (2, 26) = 26.47, P < 0.0001. CT vs. aSR (P = 0.0288), CT vs. aSS (P = 0.0002). **(E)** F (2, 23) = 0.09846, P = 0.9066. CT vs. aSR (P = 0.9966), CT vs. aSS (P = 0.9091). **(F)** F (2, 28) = 0.6065, P = 0.5522. CT vs. aSR (P = 0.4717), CT vs. aSS (P = 0.9396). **(H)** F (4, 46) = 5.085, P = 0.0018. CT vs. 0 h (P = 0.0009), CT vs. 0.5 h (P = 0.0062), CT vs. 1 h (P = 0.1531), CT vs. 2 h (P = 0.4515). **(I)** F (4, 45) = 4.309, P = 0.0049. CT vs. 0 h (P = 0.0096), CT vs. 0.5 h (P = 0.0436), CT vs. 1 h (P = 0.9998), CT vs. 2 h (P = 0.8939). **(K)** Test1: F (2, 45) = 26.92, P < 0.0001, CT vs. aSR-aSS (P = 0.0048), CT vs. aSS-aSR (P = 0.0003). Test2: F (2, 45) = 16.09, P < 0.0001, CT vs. aSR-aSS (P = 0.0066), CT vs. aSS-aSR (P = 0.0249).

**Figure 2:**

**(C)** t = 16.63, df = 17, P < 0.0001. **(D)** Column factor: F (2, 70) = 8.471, P = 0.0005. Row factor: F (1, 70) = 6.139, P = 0.0156. Interaction: F (2, 70) = 9.941, P = 0.0002. Sham CT vs. Sham aSR (P = 0.0250), Sham CT vs. Sham aSS (P = 0.0363), Sham CT vs. X-ray CT (P = 0.9809), Sham CT vs. X-ray aSR (P = 0.5658), Sham CT vs. X-ray aSS (P = 0.4027), X-ray CT vs. X-ray aSR (P = 0.9411), X-ray CT vs. X-ray aSS (P = 0.8432). **(G)** t = 6.398, df = 38, P < 0.0001. **(H)** Column factor: F (2, 83) = 15.31, P < 0.0001. Row factor: F (1, 83) = 3.402, P = 0.0687. Interaction: F (2, 83) = 9.575, P = 0.0002. Rov-EGFP CT vs. Rov-EGFP aSR (P = 0.0406), Rov-EGFP CT vs. Rov-EGFP aSS (P = 0.0016), Rov-EGFP CT vs. Rov-Cre CT (P = 0.9914), Rov-EGFP CT vs. Rov-Cre aSR (P = 0.8369), Rov-EGFP CT vs. Rov-Cre aSS (P = 0.9991), Rov-Cre CT vs. Rov-Cre aSR (P = 0.9895), Rov-Cre CT vs. Rov-Cre aSS (P > 0.9999). **(K)** t = 10.29, df =10, P < 0.0001. **(L)** Column factor: F (2, 64) = 17.57, P < 0.0001. Row factor: F (1, 64) = 2.224, P = 0.1408. Interaction: F (2, 64) = 9.814, P = 0.0002. Young CT vs. Young aSR (P = 0.0208), Young CT vs. Young aSS (P = 0.0010), Young CT vs. Aged CT (P = 0.8398), Young CT vs. Aged aSR (P = 0.9853), Young CT vs. Aged aSS (P = 0.9992), Aged CT vs. Aged aSR (P = 0.9958), Aged CT vs. Aged aSS (P = 0.6756).

**Figure 3:**

**(C)** F (2, 13) = 12.06, P = 0.0011. CT vs. aSR (P = 0.0008), CT vs. aSS (P = 0.0064). **(G)** Column factor: F (2, 85) = 11.61, P < 0.0001. Row factor: F (1, 85) = 0.7819, P = 0.3790. Interaction: F (2, 85) = 9.887, P = 0.0001. Rov-EGFP CT vs. Rov-EGFP aSR (P = 0.0051), Rov-EGFP CT vs. Rov-EGFP aSS (P = 0.0087), Rov-EGFP CT vs. Rov-Cre CT (P = 0.9762), Rov-EGFP CT vs. Rov-Cre aSR (P = 0.9902), Rov-EGFP CT vs. Rov-Cre aSS (P = 0.9993), Rov-Cre CT vs. Rov-Cre aSR (P > 0.9999), Rov-Cre CT vs. Rov-Cre aSS (P = 0.9993).

**Figure 4:**

**(C)** F (2, 16) = 35.12, P < 0.0001. CT vs. aSR (P = 0.0013), CT vs. aSS (P = 0.0020). **(D)** F (2, 17) = 5.766, P = 0.0123. CT vs. aSR (P = 0.0098), CT vs. aSS (P = 0.0300). **(E)** F (2, 16) = 33.56, P < 0.0001. CT vs. aSR (P = 0.2474), CT vs. aSS (P = 0.0001). **(H)** F (2, 14) = 0.04615, P = 0.9550. CT vs. aSR (P = 0.9691), CT vs. aSS (P = 0.9960). **(I)** F (2, 14) = 2.003, P = 0.1718. CT vs. aSR (P = 0.7847), CT vs. aSS (P = 0.1308).

**Figure 5:**

**(B)** Column factor: F (1, 60) = 4.528, P = 0.0375. Row factor: F (1, 60) = 0.3521, P = 0.5552. Interaction: F (1, 60) = 6.315, P = 0.0147. Rov-EGFP CT vs. Rov-EGFP aSR (P = 0.0088), Rov-EGFP CT vs. Rov-Cre CT (P = 0.6800), Rov-EGFP CT vs. Rov-Cre aSR (P = 0.8638), Rov-Cre CT vs. Rov-Cre aSR (P > 0.9999). **(C)** Column factor: F (1, 34) = 41.99, P < 0.0001. Row factor: F (1, 34) = 2.604, P = 0.1158. Interaction: F (1, 34) = 2.005, P = 0.1659. Rov-EGFP CT vs. Rov-EGFP aSS (P = 0.0048), Rov-EGFP CT vs. Rov-Cre CT (P > 0.9999), Rov-EGFP CT vs. Rov-Cre aSS (P < 0.0001), Rov-Cre CT vs. Rov-Cre aSS (P < 0.0001). **(H)** t = 5.603, df = 7, P = 0.0008. **(K)** t = 10.24, df = 13, P < 0.0001. **(N)** t = 1.795, df = 7, P = 0.1158. **(P)** t = 0.6290, df = 12, P = 0.5412.

**Figure 6:**

**(C)** t = 4.059, df = 18, P = 0.0007. **(D)** t = 0.5711, df = 13, P = 0.5777. **(G)** t = 12.58, df = 18, P < 0.0001. **(H)** t = 0.8699, df = 10, P = 0.4048. **(J)** r = −0.5057, R^2^ = 0.2558, P < 0.0001. **(K)** Group A: r = −0.6717, R^2^ = 0.4512, P = 0.0119; Group B: r = −0.6067, R^2^ = 0.3680, P = 0.0076; Group C: r = −0.4178, R^2^ = 0.1745, P = 0.1766; Group D: r = −0.1318, R^2^ = 0.01736, P = 0.6831.

**Figure 7:**

**(B)** t = 2.362, df = 18, P = 0.0296. **(C)** t = 2.296, df = 16, P = 0.0356. **(D)** F (2, 28) = 11.68, P = 0.0002. CT vs. aSR (P = 0.8844), CT vs. aSS (P = 0.0010). **(F)** t = 2.072, df = 5, P = 0.0930. **(H)** t = 4.888, df = 16, P = 0.0002. **(I)** t = 2.327, df = 16, P = 0.0334. **(J)** F (2, 27) = 7.978, P = 0.0019. CT vs. aSR (P = 0.8950), CT vs. aSS (P = 0.0059). **(L)** t = 2.129, df = 4, P = 0.1003. **(N)** t = 5.794, df = 34, P < 0.0001. **(O)** t = 2.362, df = 17, P = 0.0304. **(Q)** t = 4.171, df = 37, P = 0.0002. **(R)** t = 2.363, df = 16, P = 0.0311.

**Figure S1:**

**(B)** t = 3.107, df = 18, P = 0.0061. **(D)** t = 2.492, df = 18, P = 0.0227.

**Figure S2:**

**(A)** t = 1.414, df = 18, P = 0.1744. **(B)** t = 0.2510, df = 18, P = 0.8047. **(C)** t = 1.306, df =18, P = 0.2080. **(D)** t = 0.3261, df = 18, P = 0.7481. **(E)** Column factor: F (1, 96) = 0.2585, P = 0.6123. Row factor: F (5, 96) = 74.11, P < 0.0001. Interaction: F (5, 96) = 0.6992, P = 0.6254. **(F)** Column factor: F (1, 108) = 0.05283, P = 0.8186. Row factor: F (5, 108) = 69.78, P < 0.0001. Interaction: F (5, 108) = 0.5847, P = 0.7116. **(G)** Column factor: F (1, 108) = 0.7959, P = 0.3743. Row factor: F (5, 108) = 93.16, P < 0.0001. Interaction: F (5, 108) = 0.2882, P = 0.9186. **(H)** Column factor: F (1, 108) = 0.2249, P = 0.6363. Row factor: F (5, 108) = 83.94, P < 0.0001. Interaction: F (5, 108) = 0.3845, P = 0.8585. **(I)** t = 0.3265, df = 18, P = 0.7478. **(J)** t=0.7487 df=17, P = 0.4643. **(K)** t = 0.7422, df = 18, P = 0.4675. **(L)** t = 0.5325, df = 17, P =0.6013.

**Figure S3:**

**(C)** t = 4.486, df = 12, P = 0.0007. **(D)** Column factor: F (2, 65) = 16.29, P < 0.0001. Row factor: F (1, 65) = 3.250, P = 0.0760. Interaction: F (2, 65) = 6.495, P = 0.0027. Vehicle CT vs. Vehicle aSR (P = 0.0138), Vehicle CT vs. Vehicle aSS (P = 0.0044), Vehicle CT vs. TMZ CT (P = 0.9413), Vehicle CT vs. TMZ aSR (P = 0.9956), Vehicle CT vs. TMZ aSS (P = 0.3546), TMZ CT vs. TMZ aSR (P = 0.9990), TMZ CT vs. TMZ aSS (P = 0.9153).

**Figure S4:**

**(C)** t = 0.009843, df = 10, P = 0.9923.

**Figure S5:**

**(D)** t = 4.580, df = 5, P = 0.0059.

**Figure S6:**

**(B)** t = 0.4312, df = 10, P = 0.6755. **(D)** t = 0.7079, df = 9, P = 0.4969. **(F)** t = 0.1972, df = 8, P = 0.8486.

**Figure S7:**

**(D)** F (2, 20) = 36.94, P < 0.0001. Dox on vs. Dox off 2 day without training (P = 0.3967), Dox on vs. Dox off 2 day after training 4h (P < 0.0001), Dox off 2 day without training vs. Dox off 2 day after training 4h (P < 0.0001).

**Figure S8:**

**(B)** t= 0.2029, df = 14, P = 0.8422.

**Figure S9:**

**(B)** t = 0.3807, df = 10, P = 0.7114. **(D)** t = 0.2726, df = 8, P = 0.7920.

**Figure S10:**

**(C)** t = 4.699, df = 13, P = 0.0004. **<u>(D)</u>** Column factor: F (2, 90) = 55.13, P < 0.0001. Row factor: F (1, 90) = 0.02334, P = 0.8789. Interaction: F (2, 90) = 0.5318, P = 0.5894. Sedentary CT vs. Sedentary aSR (P = 0.0356), Sedentary CT vs. Sedentary aSS (P < 0.0001), Sedentary CT vs. VR CT (P = 0.9696), VR CT vs. VR aSR (P = 0.0101), VR CT vs. VR aSS (P = 0.0160).

## Supplemental figures

**Figure S1.**
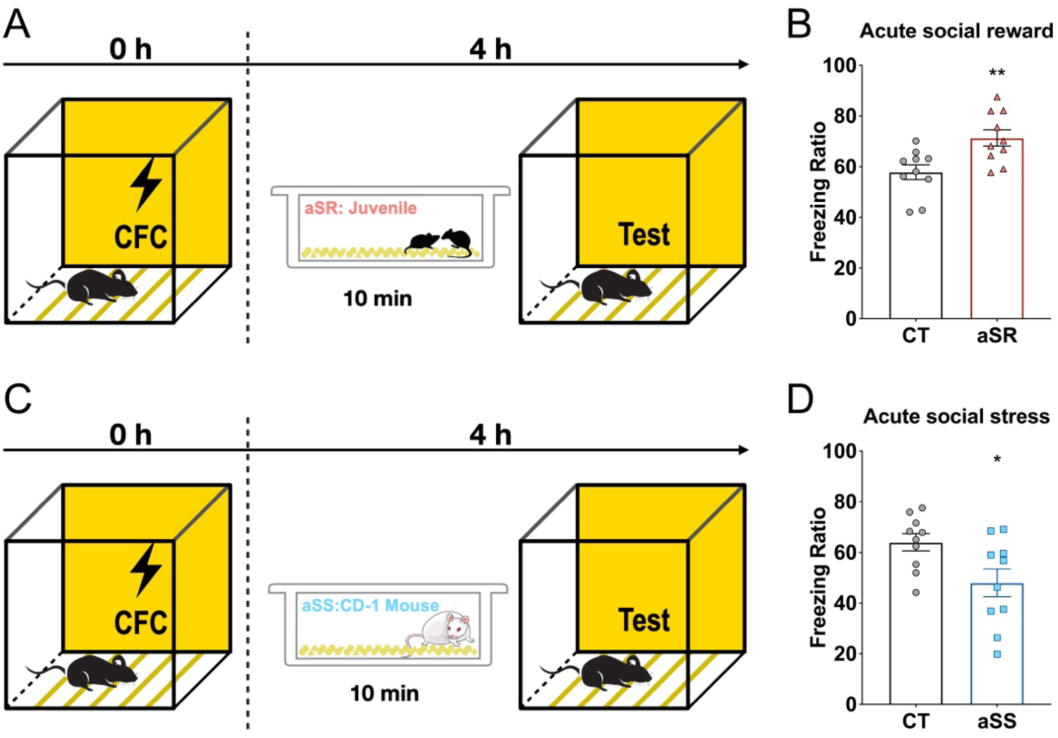
Bidirectional regulation of retrieval through aSR induced by interacting with juvenile and aSS induced by social defeat. **(A and C)** Experimental schedule of acute social experiences before retrieval. **(B)** 4 h memory with aSR induced by interacting with juvenile (n_CT_=10, n_aSR_=10). **(D)** 4 h memory with aSS induced by social defeat (n_CT_=10, n_aSS_=10). *P < 0.05, **P < 0.01; unpaired two-tailed t test. Graphs show means ± SEM.

**Figure S2.**
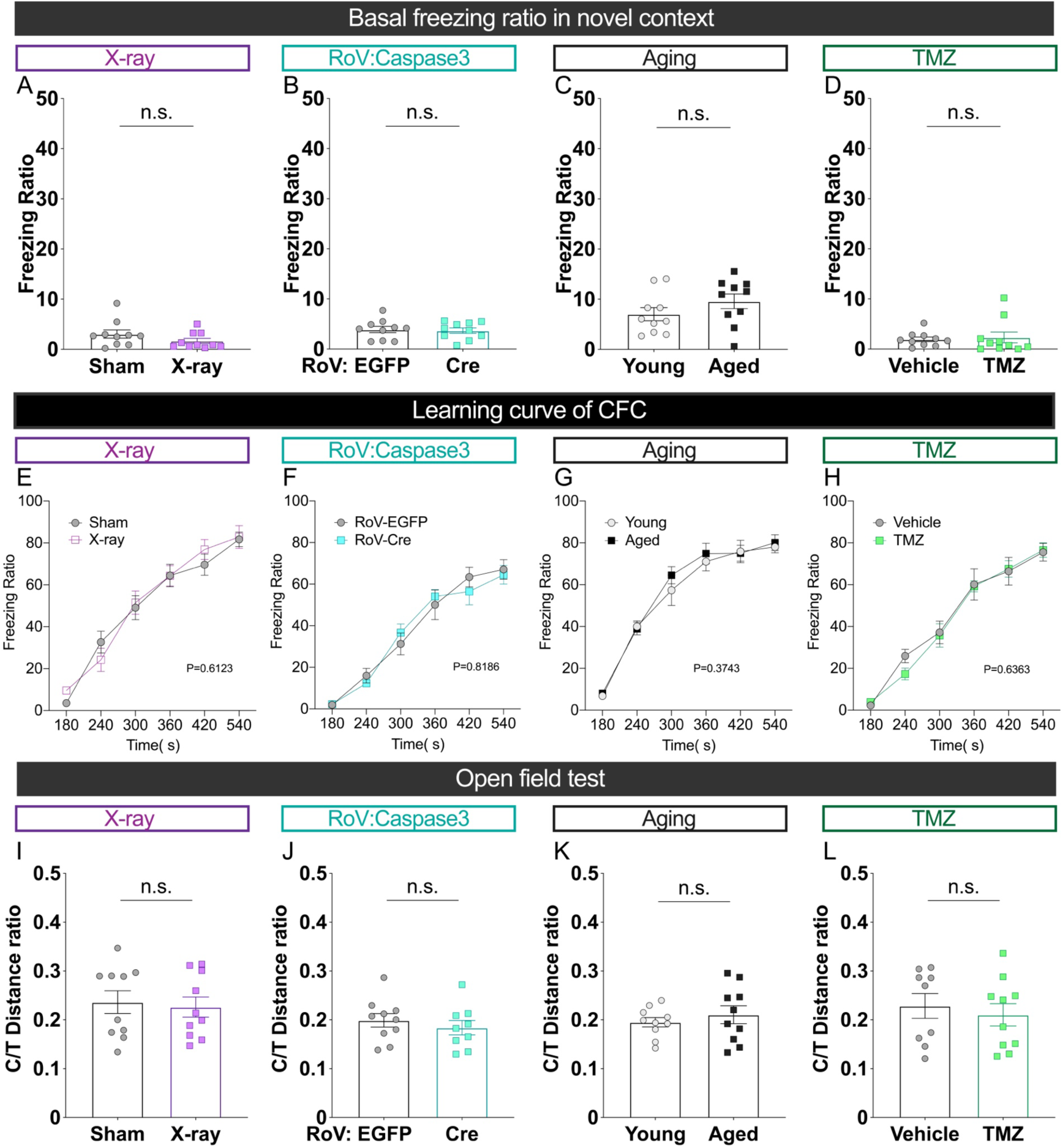
Abolishing adult hippocampal neurogenesis does not affect basal freezing, five-shock contextual fear memory acquisition and anxiety-like behavior. **(A-D)** Abolishing adult hippocampal neurogenesis does not affect basal freezing level (A, n_sham_=10, n_x-ray_=10; B, n_Rov:EGFP_=10, n_cre_=10; C, n_young_=10, n_aged_=10; D, n_vehicle_=10, n_TMZ_=10). **(E-H)** Abolishing adult hippocampal neurogenesis does not affect learning curve of contextual fear memory (E, n_sham_=9, n_x-ray_=9; F, n_Rov:EGFP_=10, n_cre_=10; G, n_young_=10, n_aged_=10; H, n_vehicle_=10, n_TMZ_=10). **(I-L)** Abolishing adult hippocampal neurogenesis does not affect performance in OFT (I, n_sham_=10, n_x-ray_=10; J, n_Rov:EGFP_=10, n_cre_=9; K, n_young_=10, n_aged_=10; L, n_vehicle_=9, n_TMZ_=10). Unpaired two-tailed t test (A-D and I-L) or two-way analysis of variance (ANOVA) (E-H). Graphs show means ± SEM.

**Figure S3.**
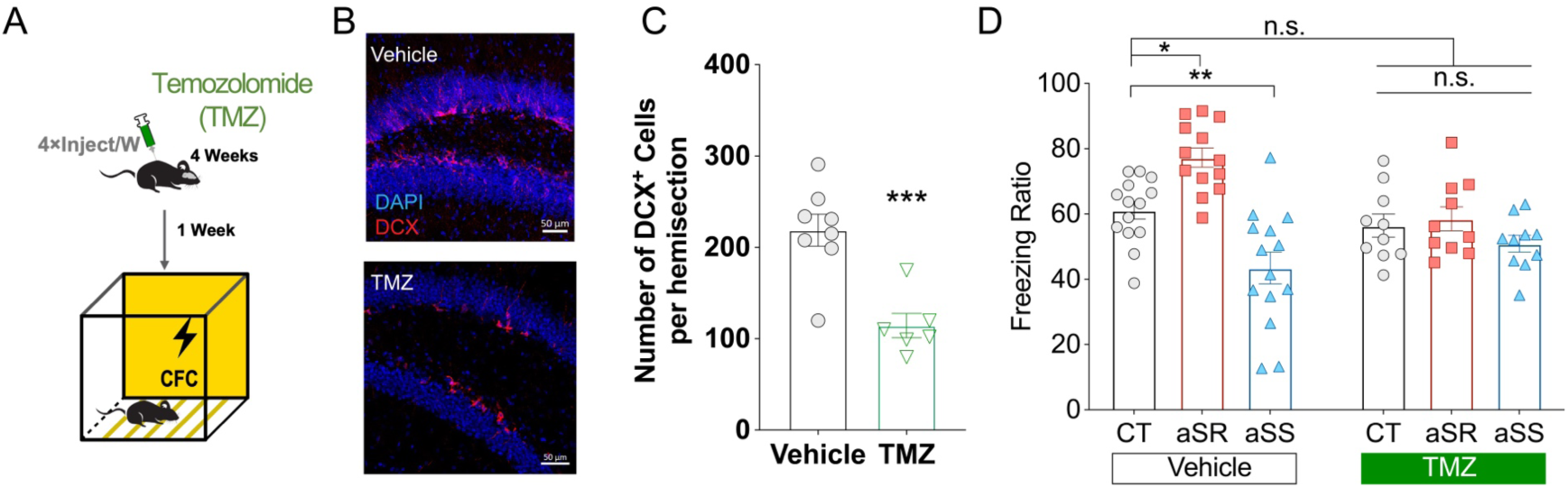
Temozolomide treatment decreases neurogenesis and blocks the effects of acute social experiences. **(A)** Experimental schedule of anbGC ablation through temozolomide (TMZ) injection. **(B)** Representative images of anbGCs (DCX) in the dDG. **(C)** Quantification of anbGCs in the dDG (n_Vehicle_=8, n_TMZ_=6). **(D)** 4 h memory in the vehicle-treated group (n_CT_=14, n_aSR_=13, n_aSS_=14) and in the TMZ-treated group (n_CT_=10, n_aSR_=10, n_aSS_=10). *P < 0.05, **P < 0.01, ***P < 0.001; unpaired two-tailed t test (C), two-way analysis of variance (ANOVA) with Tukey’s test (D). Graphs show means ± SEM.

**Figure S4.**
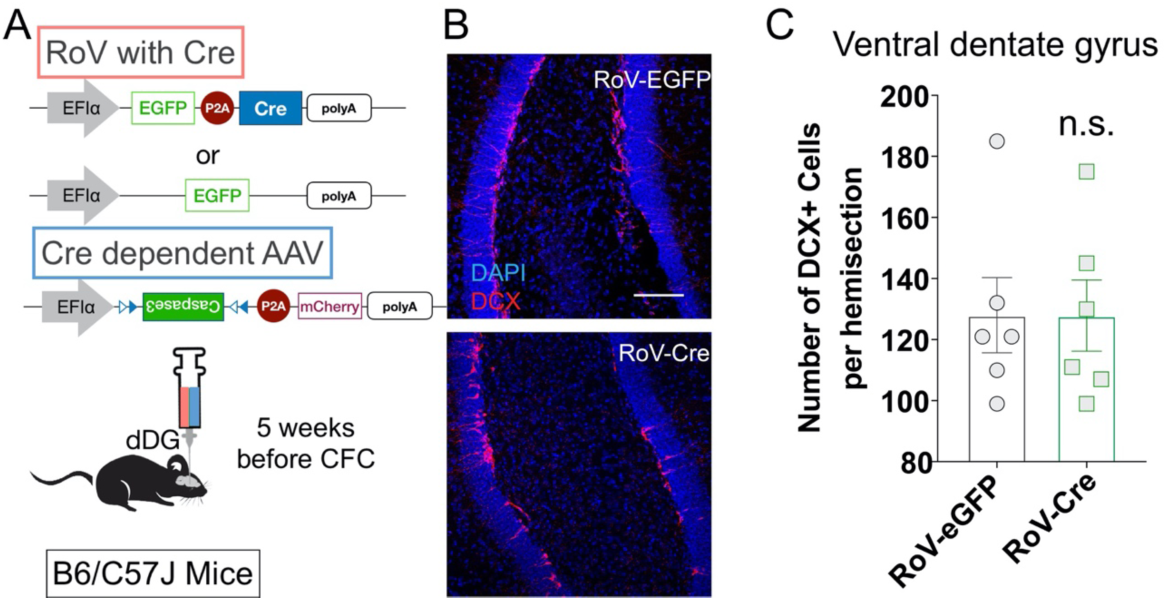
Double virus-mediated anbGC ablation in dDG does not influence anbGCs in ventral dentate gyrus (vDG). **(A)** Experimental schedule of anbGC ablation in dDG through caspase3 expression driven by double-virus system. **(B)** Representative images of anbGCs (DCX) in the vDG, scale bar: 50μm. **(C)** Quantification of anbGCs in the vDG (n_RoV-eGFP_=6, n_RoV-Cre_=6). Unpaired two-tailed t test. Graphs show means ± SEM.

**Figure S5.**
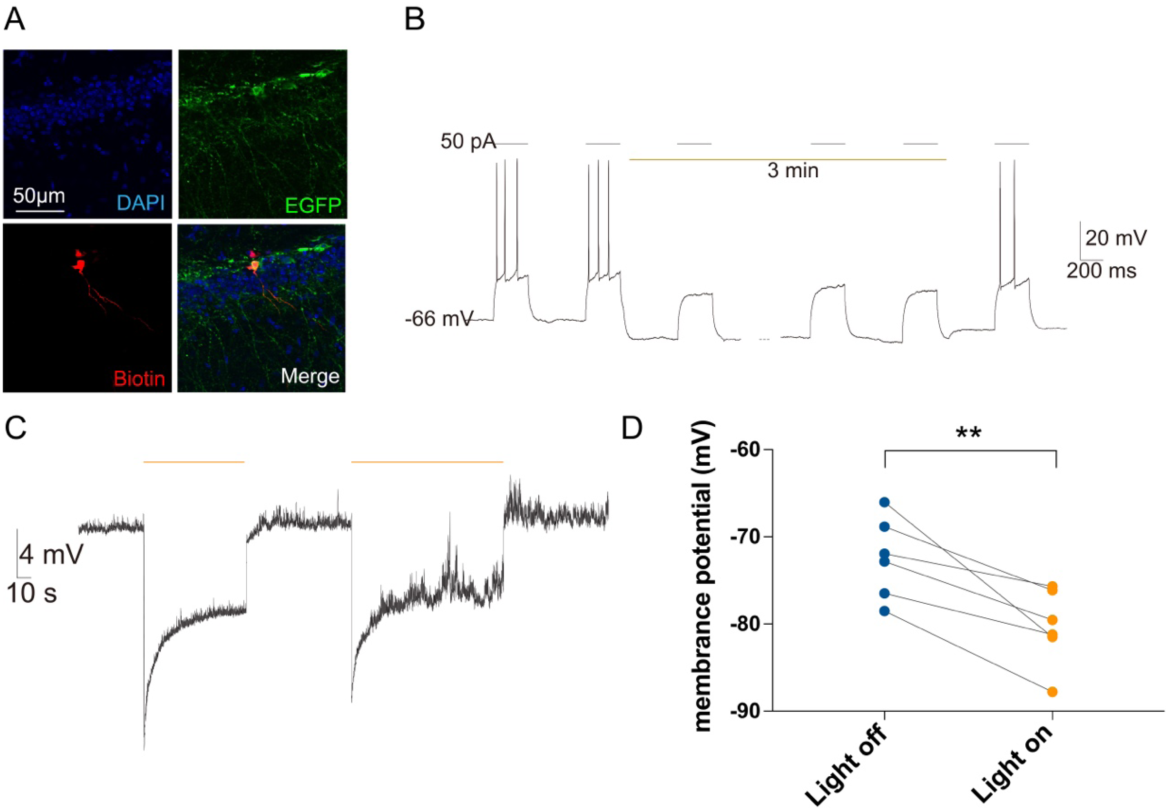
Functional verification of expressing eNpHR3.0 in dDG. **(A)** Representative images showing that biotin was injected into EGFP^+^ cells during the in vitro whole-cell recording. **(B)** Electrophysiological recording of anbGCs with persistent optogenetical inhibition. **(C)** Representative trace of resting potential in anbGCs upon 590 nm laser stimulation. **(D)** 590 nm laser decreased the resting membrane potential of anbGCs (n=6). **P < 0.01; paired two-tailed t test. Graphs show means ± SEM.

**Figure S6.**
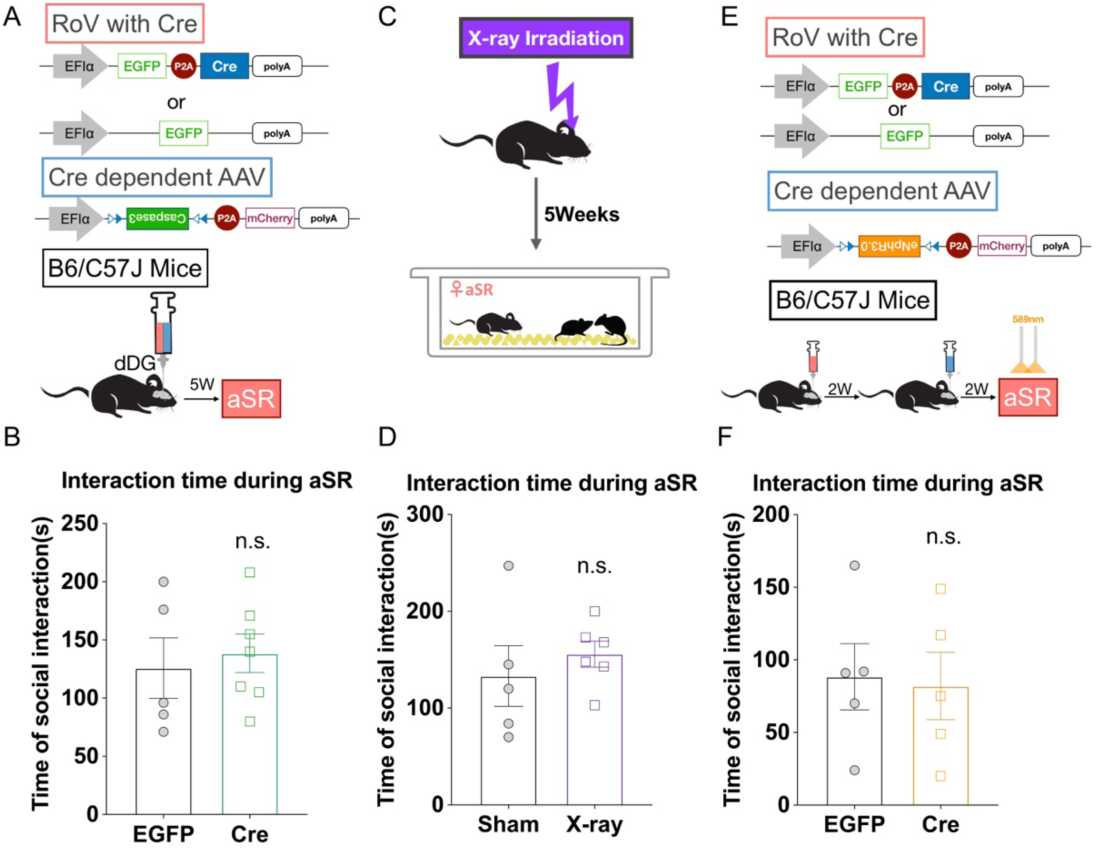
Unaltered social interaction time during aSR in mice with ablation or inhibition of anbGCs. **(A and C)** Experimental schedules for anbGC ablation. **(B and D)** Social interaction time during aSR in mice with anbGC ablation by double-virus system (B, n_EGFP_=5, n_Cre_=7) or by X-ray irradiation (D, n_Sham_=5, n_X-ray_=6). **(E)** Experimental schedule for anbGC inhibition during aSR. **(F)** Social interaction time during aSR in mice with anbGC inhibition during aSR (n_EGFP_=5, n_Cre_=5). Unpaired two-tailed t test. Graphs show means ± SEM.

**Figure S7.**
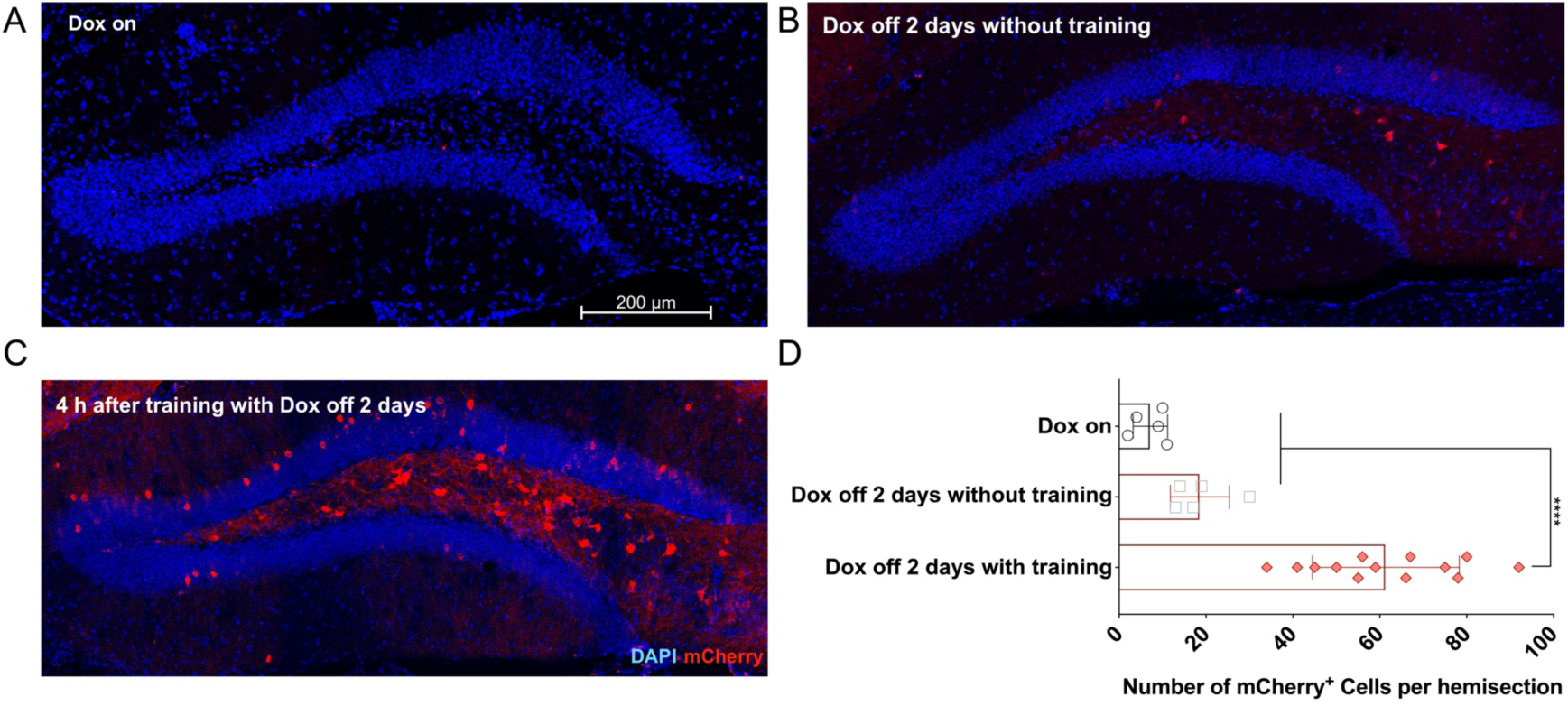
The efficiency of activity-dependent expression of mCherry in the dDG by AAV during CFC. **(A-C)** Representative images of the dDG from mice injected with AAV-c-Fos-tTA and AAV-TRE-mCherry that were euthanised after the following treatments: Dox on (A); Dox off for 2 days without training (B); Dox off for 2 days, followed by CFC training and perfused 4 h after training (C). **(D)** Quantification of mCherry^+^ cells (n_Dox on_=5, n_Dox off 2 days without training_=5, n_Dox off 2 day after training 4h_=13). ****P < 0.0001; one-way ANOVA with Dunnett’s test. Graphs show means ± SEM.

**Figure S8.**
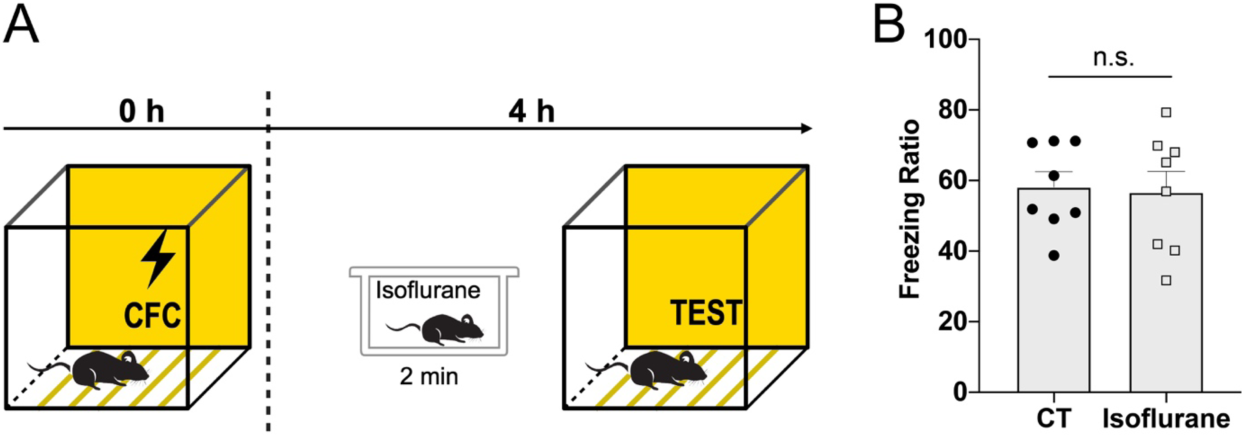
Isoflurane-induced anesthesia does not influence 5-shock contextual fear memory retrieval. **(A)** Experimental schedule. **(B)** 4 h memory with isoflurane-induced anesthesia before retrieval, (n_CT_=8, n_isoflurane_ =8). Unpaired t-test. Graphs show means ± SEM.

**Figure S9.**
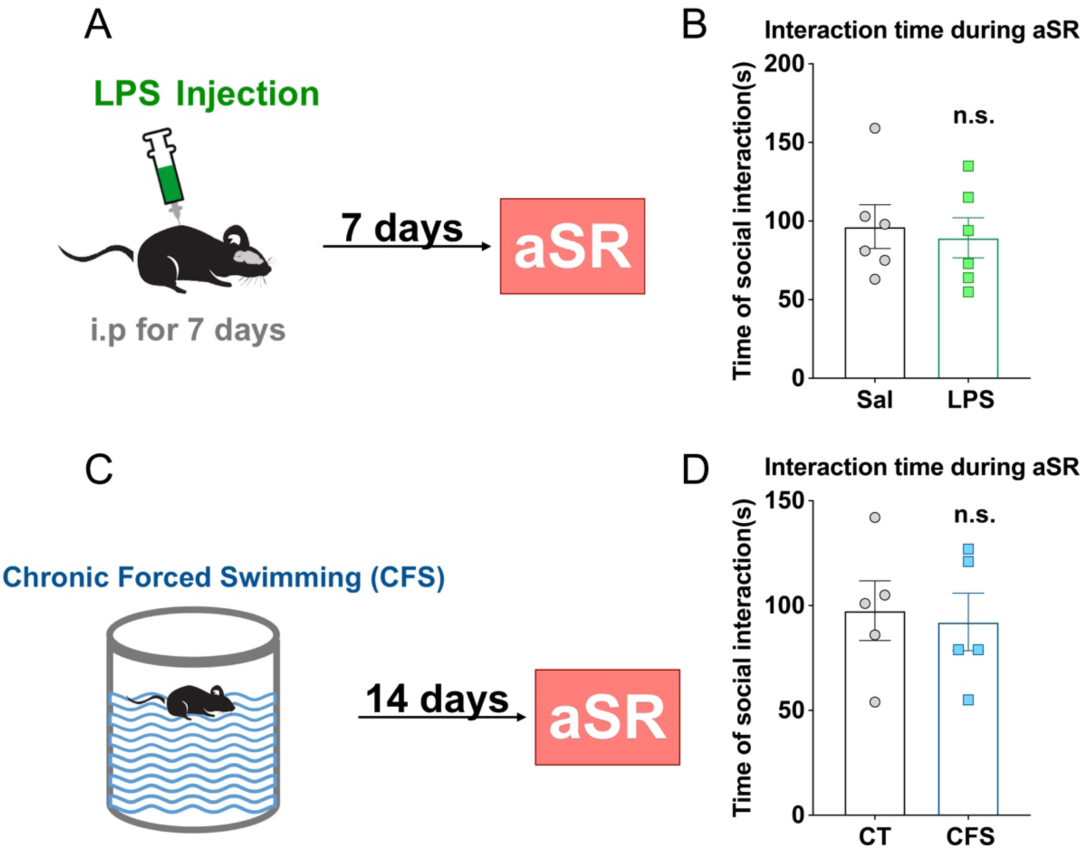
Unaltered social interaction time during aSR in stress models. **(A and C)** Stress models induced in mice by LPS injection (A) and by chronic forced swimming (C), and experimental schedules. **(B and D)** Social interaction time during aSR in stress models (B: n_Sal_=6, n_LPS_=6; D: n_CT_=5, n_CFS_=5). Unpaired two-tailed t test. Graphs show means ± SEM.

**Figure S10.**
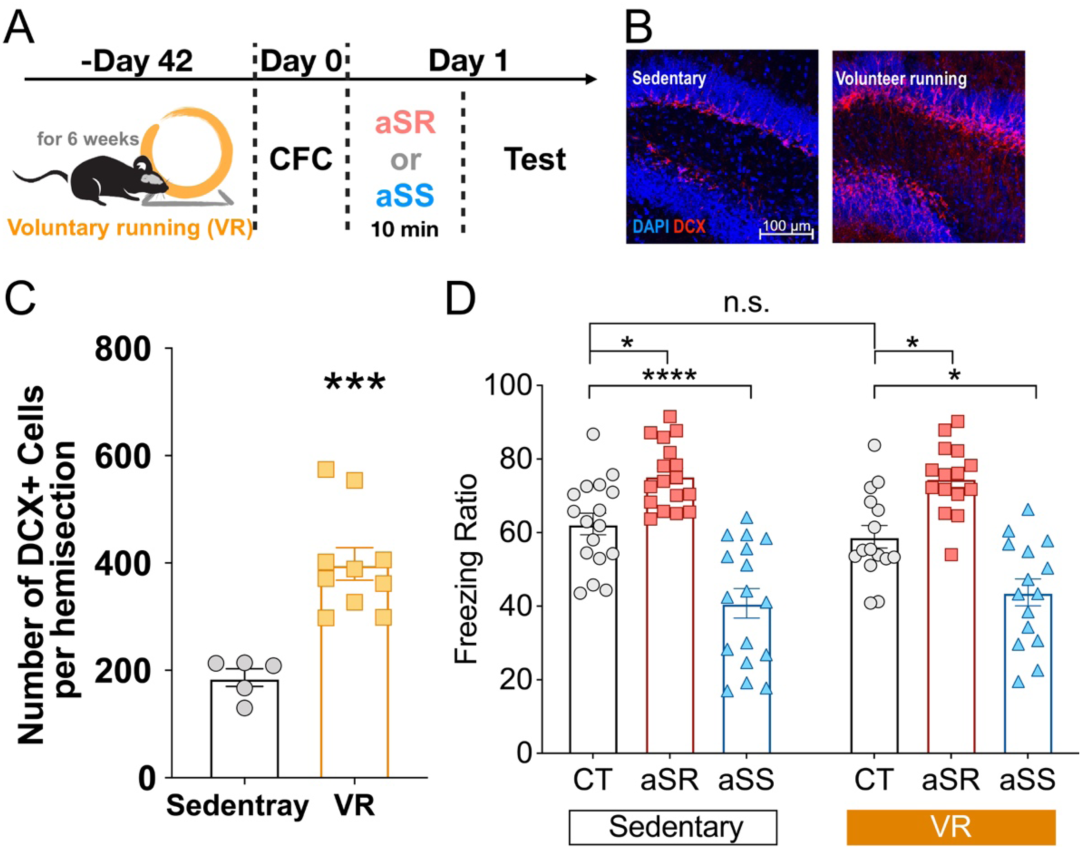
Voluntary running can increase neurogenesis without affecting the effect of acute social experiences. **(A)** Experimental schedule. **(B)** Representative images of anbGCs (DCX) in the dDG. **(C)** Quantification of anbGCs in the dDG (n_Sedentary_=5, n_VR_=10). **(D)** 4 h memory with acute social experiences in sedentary mice (n_CT_=17, n_aSR_=17, n_aSS_=17) and with acute social experiences in mice with voluntary running (n_CT_=15, n_aSR_=15, n_aSS_=15). *P < 0.05, ***P < 0.001, ****P < 0.0001; unpaired two-tailed t test (C) or two-way analysis of variance (ANOVA) with Tukey’s test (D). Graphs show means ± SEM.

